# Altered structural brain asymmetry in autism spectrum disorder: large-scale analysis via the ENIGMA Consortium

**DOI:** 10.1101/570655

**Authors:** Merel C. Postema, Daan van Rooij, Evdokia Anagnostou, Celso Arango, Guillaume Auzias, Marlene Behrmann, Geraldo Busatto Filho, Sara Calderoni, Rossa Calvo, Eileen Daly, Christine Deruelle, Adriana Di Martino, Ilan Dinstein, Fabio Luis S. Duran, Sarah Durston, Christine Ecker, Stefan Ehrlich, Damien Fair, Jennifer Fedor, Xin Feng, Jackie Fitzgerald, Dorothea L. Floris, Christine M. Freitag, Louise Gallagher, David C. Glahn, Ilaria Gori, Shlomi Haar, Liesbeth Hoekstra, Neda Jahanshad, Neda Jahanshad, Maria Jalbrzikowski, Joost Janssen, Joseph A. King, Luisa L Zaro, Jason P. Lerch, Beatriz Luna, Mauricio M. Martinho, Jane McGrath, Sarah E. Medland, Filippo Muratori, Clodagh M. Murphy, Declan G.M. Murphy, Kirsten O’Hearn, Bob Oranje, Mara Parellada, Olga Puig, Alessandra Retico, Pedro Rosa, Katya Rubia, Devon Shook, Margot Taylor, Michela Tosetti, Gregory L. Wallace, Fengfeng Zhou, Paul Thompson, Simon E. Fisher, Jan K. Buitelaar, Clyde Francks

**Affiliations:** Department of Language & Genetics, Max Planck Institute for Psycholinguistics, Nijmegen, the Netherlands; Department of Cognitive Neuroscience, Donders Institute for Brain, Cognition and Behaviour, Donders Centre for Cognitive Neuroimaging, Radboud University Medical Centre, Nijmegen, The Netherlands; Bloorview Research Institute, Holland Bloorview Kids Rehabilitation Hospital and Department of pediatrics, University of Toronto, Toronto, Canada, USA; Child and Adolescent Psychiatry Department, Gregorio Marañón General University Hospital, School of Medicine, Universidad Complutense, IiSGM, CIBERSAM, Madrid, Spain; Institut de Neurosciences de la Timone, UMR 7289, Aix Marseille Université, CNRS, Marseille, France; Department of Psychology, Carnegie Mellon University, Pittsburgh, PA, USA; Laboratory of Psychiatric Neuroimaging (LIM-21), Departamento e Instituto de Psiquiatria, Hospital das Clinicas HCFMUSP, Faculdade de Medicina, Universidade de Sao Paulo, Sao Paulo, SP, BR; IRCCS Stella Maris Foundation, viale del Tirreno 331, 56128, Pisa, Italy; Department of Child and Adolescent Psychiatry and Psychology Hospital Clinic, Barcelona CIBERSAM, Universitat de Barcelona; Department of Forensic and Neurodevelopmental Sciences, Institute of Psychiatry, Psychology & Neuroscience King’s College London, London, UK.; Institute for Pediatric Neuroscience, NYU Child Study Center, NY, USA; Department of Psychology, Ben-Gurion University of the Negev, Beer Sheva, Israel; Brain Center Rudolf Magnus, Department of Psychiatry, University Medical Center Utrecht, The Netherlands; Department of Child and Adolescent Psychiatry, Psychosomatics and Psychotherapy, University Hospital, Goethe University Frankfurt am Main, Frankfurt, Germany; Division of Psychological and Social Medicine and Developmental Neurosciences, Faculty of Medicine, TU Dresden, Germany; Department of Behavioral Neuroscience, Oregon Health & Science University, Portland, Oregon, USA; Department of Psychiatry, University of Pittsburgh, Pittsburgh, PA, USA; Department of Psychiatry, School of Medicine, Trinity College, Dublin, Ireland; Department of Psychiatry, Boston Children’s Hospital and Harvard Medical School, Boston, MA 02115-5724, USA; National Institute for Nuclear Physics, Pisa Division, Largo B. Pontecorvo 3, 56124, Pisa, Italy; Department of Bioengineering, Imperial College London, London, United Kingdom; University of Sourthern California, Imaging Genetics Center, Mark & Mary Stevens Institution for Neuroimaging & Informatics; Imaging Genetics Center, Mark and Mary Stevens Neuroimaging and Informatics Institute, Keck School of Medicine of USC, Marina del Rey, CA 90292 USA; Department of Child and Adolescent Psychiatry and Psychology Hospital Clinic, Barcelona CIBERSAM, Universitat de Barcelona, IDIBAPS; Neurosciences and Mental Health, The Hospital for Sick Children, Toronto, Canada, USA; Department of Neuropsychiatry, Universidade Federal de Santa Maria, Brazil; Psychiatric Genetics, QIMR Berghofer Medical Research Institute, Brisbane, Queensland, Australia; The Sackler Institute for Translational Neurodevelopment, Institute of Psychiatry, Psychology & Neuroscience, King’s College London, London, UK; Department of Child and Adolescent Psychiatry and Psychology Hospital Clinic, Barcelona, 2017SGR881. CIBERSAM, Spain.; Institute of Psychiatry, Psychology and Neuroscience, Kings College London, London, UK; Diagnostic Imaging Research, The Hospital for Sick Children, University of Toronto, Canada; Department of Speech and Hearing Sciences, The George Washington University, Washington, DC, USA; BioKnow Health Informatics Lab, College of Computer Science and Technology, and Key Laboratory of Symbolic Computation and Knowledge Engineering of Ministry of Education, Jilin University, Changchun, Jilin, 130012, China.; Department of Clinical and Experimental Medicine, University of Pisa, Pisa, Italy; Imaging Genetics Center, Mark & Mary Stevens Institution for Neuroimaging & Informatics, University of Southern California, Marina del Rey, CA, USA; Department of Child and Adolescent Psychiatry, Division of Psychological and Social Medicine and Developmental Neurosciences, Faculty of Medicine, TU Dresden, Germany; The Trinity College Institute of Neuroscience, Trinity College, Dublin, Ireland; Olin Neuropsychiatric Research Center, Hartford, CT, USA; Karakter Child and Adolescent Psychiatry University Centre, Nijmegen, The Netherlands; Department of Medical Biophysics, University of Toronto, Toronto, Canada, USA.; Behavioural Genetics Clinic, Adult Autism Service, Behavioural and Developmental Psychiatry Clinical Academic Group, South London and Maudsley Foundation NHS Trust, London, UK; Donders Institute for Brain, Cognition and Behaviour, Radboud University, Nijmegen, The Netherlands

**Author notes:** Corresponding author: Clyde Francks, DPhil, Max Planck Institute for Psycholinguistics, Wundtlaan 1, Nijmegen, The Netherlands.

**Keywords:** autism, brain asymmetry, brain laterality, mega-analysis, structural imaging, cortical thickness

## Abstract

**Background:** Left-right asymmetry is an important organizing feature of the healthy brain. Various studies have reported altered structural brain asymmetry in autism spectrum disorder (ASD). However, findings have been inconsistent, likely due to limited sample sizes and low statistical power.

**Methods:** We investigated 1,774 subjects with ASD and 1,809 controls, from 54 datasets, for differences in the asymmetry of thickness and surface area of 34 cerebral cortical regions. We also examined global hemispheric measures of cortical thickness and area asymmetry, and volumetric asymmetries of subcortical structures. Data were obtained via the ASD Working Group of the ENIGMA (Enhancing NeuroImaging Genetics through Meta-Analysis) consortium. T1-weighted MRI data were processed with a single protocol using FreeSurfer and the Desikan-Killiany atlas.

**Results:** ASD was significantly associated with reduced leftward asymmetry of total hemispheric average cortical thickness, compared to controls. Eight regional thickness asymmetries, distributed over the cortex, also showed significant associations with diagnosis after correction for multiple comparisons, for which asymmetry was again generally lower in ASD versus controls. In addition, the medial orbitofrontal surface area was less rightward asymmetric in ASD than controls, and the putamen volume was more leftward asymmetric in ASD than controls. The largest effect size had Cohen’s *d* = 0.15. Most effects did not depend on age, sex, IQ, or disorder severity.

**Conclusion:** Altered lateralized neurodevelopment is suggested in ASD, affecting widespread cortical regions with diverse functions. Large-scale analysis was necessary to reliably detect, and accurately describe, subtle alterations of structural brain asymmetry in this disorder.

## Introduction

Autism spectrum disorder (ASD) is an umbrella diagnosis, capturing several previously separate pervasive developmental disorders with various levels of symptom severity, including Autistic Disorder, Asperger’s Syndrome, Childhood Disintegrative Disorder, and Pervasive Developmental Disorder – Not Otherwise Specified (PDD-NOS) (1). According to the Diagnostic and Statistical Manual of Mental Disorders (DSM) version 5, diagnosis of ASD requires the presence of at least three symptoms of impaired social communication and at least two symptoms of repetitive behaviours or restricted interests (1). ASD has a prevalence of 2-3% in United States children (2).

Characterizing the neurobiology of ASD may eventually lead to improved diagnosis and clinical subgrouping, and the development of individually targeted treatment programs (3). While much of the neurobiology of ASD remains unknown, subtle alterations of brain structure appear to be involved (reviewed by (4, 5)). These include differences in total brain volume (children with ASD have shown a larger average volume), as well as alterations in the inferior frontal, superior temporal, and orbitofrontal cortices, and the caudate nucleus. However, the results of structural MRI studies of ASD have often been inconsistent, due to 1) small study sample sizes in relation to subtle effects, 2) differences across studies in terms of clinical characteristics, age, comorbidity and medication use, 3) methodological differences between studies, such as differences in hardware, software and distinct data processing pipelines (6), and 4) the etiological and neurobiological heterogeneity of ASD, which exists as a group of different syndromes rather than a single disease entity (7).

In the ENIGMA (Enhancing Neuro-Imaging Genetics through Meta-Analysis) consortium (http://enigma.ini.usc.edu), researchers from around the world collaborate to analyse many separate datasets jointly, and to reduce some of the technical heterogeneity by using harmonized protocols for MRI data processing. A recent study by the ENIGMA consortium’s ASD working group showed small average differences in bilateral cortical and subcortical brain measures between 1,571 cases and 1,650 healthy controls, in the largest study of brain structure in ASD yet performed (8). Relative to controls, ASD patients had significantly lower volumes of several subcortical structures, as well as greater thickness in various cortical regions - mostly in the frontal lobes - and lower thickness of temporal regions. No associations of diagnosis with regional cortical surface areas were found (8).

Left-right asymmetry is an important aspect of human brain organization, which may be altered in various psychiatric and neurocognitive conditions, including schizophrenia, dyslexia and ASD (9-11). On a functional level, people with ASD demonstrate reduced leftward language lateralization more frequently than controls (12-14). Resting state functional magnetic resonance imaging (MRI) data have also shown a generally rightward shift of asymmetry involving various functional networks of brain regions (15). In addition, people with ASD have a higher rate of left-handedness than the general population (14, 16, 17). Furthermore, an electroencephalography study reported that infants at high risk for ASD showed more rightward than leftward frontal alpha asymmetry at rest (18).

Brain structural imaging studies have also reported altered hemispheric asymmetry in ASD. Diffusion imaging studies indicated reduced asymmetry of a variety of different white matter tract metrics (19-21), although in one study males with ASD lacked an age-dependent decrease in rightward asymmetry of network global efficiency, compared to controls (22). A structural MRI study investigating grey matter reported lower leftward volume asymmetry of language-related cortical regions in ASD (i.e., *planum temporale*, Hesch’s gyrus, posterior supramarginal gyrus and parietal operculum), as well as greater rightward asymmetry of the inferior parietal lobule (23). The volume and surface area of the fusiform gyrus also showed lower rightward asymmetry in ASD (24). However, other studies did not find alterations of grey matter asymmetries in ASD (21, 25).

Prior studies of structural brain asymmetry in ASD had sample sizes less than 128 cases and 127 controls. The previous ENIGMA consortium study of ASD (8) did not perform analyses of brain asymmetry, but reported bilateral effects only as strong as Cohen’s *d* = −0.21 (for the entorhinal thickness bilaterally) (8). Comparable bilateral effect sizes were also found in ENIGMA consortium studies of other disorders (8, 26-32). If effects on brain asymmetry are similarly subtle, then prior studies of this aspect of brain structure in ASD were likely underpowered. Low power not only reduces the chance of detecting true effects, but also the likelihood that a statistically significant result reflects a true effect (33, 34). Therefore a large scale analysis was needed to determine whether, and how, structural brain asymmetry might be altered in ASD, to better describe the neurobiology of the condition.

Here, we made use of MRI data from 54 datasets that were collected across the world by members of the ENIGMA consortium’s ASD Working Group, to perform the first highly-powered study of structural brain asymmetry in ASD. Using a single, harmonized protocol for image analysis, we derived asymmetry indexes, AI= (Left-Right)/(Left+Right), for multiple brain regional and global hemispheric measures, in up to 1,778 individuals with ASD and 1,829 typically developing controls. The AI is a widely used index in brain asymmetry studies (35, 36). Regional asymmetry indices that showed significant case-control differences were tested for age- or sex-specific effects, and also for correlations with IQ or disorder severity, which serve as key indicators of the clinical heterogeneity in ASD.

## Materials and Methods

### Datasets

Structural MRI data were available for 57 different datasets (**Table S1**). Three datasets comprising either cases only, or controls only, were removed in this study (**Table S1**), as our analysis model included random intercepts for ‘dataset’ (below), and diagnosis was fully confounded with dataset for these three. The remaining 54 datasets comprised 1,778 people with ASD (N = 1,504 males; median age = 13 years; range = 2 to 64 years) and 1,829 typically developing controls (N = 1,400 males; median age = 13 years; range = 2 to 64 years).

Diagnosis was based on clinical assessment according to either DSM version 4 or version 5 criteria (1). In addition, severity measures were available for a majority of cases (see **Table S2**), in the form of raw scores according to the Autism Diagnostic Observation Schedule-Generic (ADOS), a standardized instrument commonly used in autism diagnosis (37). Cases from the entire ASD spectrum were included, but most cases did not have an intellectual disability (cases: mean IQ = 104, SD = 19, min = 34, max = 149; controls: mean IQ = 112, SD = 15, min = 31, max = 149) (see **Figure S1C** for the distribution plots). The presence/absence of comorbid conditions had been recorded for 519 of the cases, but only 54 cases showed at least one comorbid condition (which could be attention-deficit hyperactivity disorder (ADHD), obsessive compulsive disorder (OCD), depression, anxiety, and/or Tourette’s syndrome (8)).

### Structural magnetic resonance imaging

Structural T1-weighted brain MRI scans were acquired at each study site. As shown in Supplementary Table S1, images were acquired using different field strengths (1.5 T or 3 T) and scanner types. Each site used harmonized protocols from the ENIGMA consortium (http://enigma.ini.usc.edu/protocols/imaging-protocols) for data processing and quality control. The data used in the current study were thickness and surface area measures for each of 34 bilaterally paired cortical regions, the latter as defined with the Desikan-Killiany atlas (38), as well as the average cortical thickness and total surface area per entire hemisphere. In addition, left and right volumes of seven bilaterally paired subcortical structures, plus the lateral ventricles, were analyzed. Cortical parcellations and subcortical segmentations were performed with the freely available and validated software FreeSurfer (versions 5.1 or 5.3) (39), using the default ‘recon-all’ pipeline. Parcellations of cortical grey matter regions were visually inspected following the standardized ENIGMA quality control protocol (http://enigma.ini.usc.edu/protocols/imaging-protocols). Exclusions on the basis of this quality control resulted in the sample sizes mentioned above (see Datasets).

Two of the datasets (i.e., FSM and MYAD) included participants as young as 1.5 years of age. As segmentation of very young brains might be especially challenging for the FreeSurfer algorithms, we repeated our main analysis (below) excluding these datasets, to check that they did not impact the findings substantially (although they had passed the same quality control procedures as all other datasets).

### Asymmetry measures

Left (L) and right (R) data for each structural measure and individual subject were loaded into R (version 3.3.3). For each L and R measure separately, outliers were defined per group (cases/controls) and per dataset as those values above and below 1.5 times the interquartile range, and were excluded. There were 159 to 464 outliers removed, depending on the specific measure. An asymmetry index (AI) was calculated for each subject and each paired L and R measure using the following formula: (*L – R*)/(*L* + *R*). Outliers were removed for each AI, following the same criteria as above (37 to 155 per AI). Primary analysis in this study was based on AIs, but post hoc analyses were also performed using the separate L and R measures for some regions(see below). To ensure that the identical set of subjects was used for analyses based on AI, L and R data, all values that were missing in the AI data were also removed from the L and R data. Distributions of each of the AIs after outlier removal are plotted in **Figure S2**.

We removed outliers separately for each AI because analyses were performed separately per AI (see below), and each AI needed to behave in a robust way in the linear model used. However, we also repeated the main analyses without removing any outliers from the L, R or AI measures, to confirm that the results were not overly dependent on outlier removal.

### Linear mixed effects model mega-analysis

#### Model

Linear mixed effects models were fitted separately for each cortical regional surface and thickness AI, as well as the total hemispheric surface area and mean thickness AI, and the subcortical volume AIs. This was accomplished by means of mega-analysis incorporating data from all 54 datasets, using the nlme package in R (40). All models included the same fixed- and random effects, and had the following formulation:

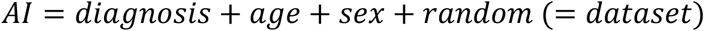

where *AI* reflects the AI of a given brain structure, and *Diagnosis* (‘controls’ (= reference), ‘ASD’), *sex* (‘males’ (= reference), ‘females’) and *dataset* were coded as factor variables, with dataset having 54 different categories. *Age* was coded as a numeric variable.

The Maximum Likelihood (ML) method was used to fit the models. Subjects were omitted if data were missing for any of the predictor variables (method = na.omit). The ggplot2 package in R was used to visualize residuals (**Figures S3-S5**). Collinearity of predictor variables was assessed using the usdm pacakge in R (version 3.3.3.).

The relationships between AIs and age showed no overt non-linearity (**Figures S6-S8**), so no polynomials for age were incorporated in the models for primary analysis. However, analyses were repeated using an additional non-linear term for age, to check whether this choice had affected the results. As Age and Age^2^ are highly correlated, we made use of the poly()-function in R for these two predictors, which created a pair of uncorrelated variables to model age effects (so-called orthogonal polynomials) (41), where one variable was linear and one non-linear.

#### Significance

Significance was assessed based on the P-values for the effects of *diagnosis* on AIs. The False Discovery Rate (FDR) (42) was estimated separately for the 35 cortical surface area AIs (i.e., 34 regional AIs and one hemispheric total AI) and the 35 cortical thickness AIs, and again for the seven subcortical structures plus lateral ventricles, each time with a FDR threshold of 0.05. Correlations between AI measures were calculated using Pearson’s R and visualized using the “corrplot” package in R (**Figures S9-S11**). Most pairwise correlations between AIs were low, with only 21 pairwise correlations either lower than −0.3 or greater than 0.3, with a minimum R = −0.362 between the caudal anterior cingulate surface area AI and superior frontal surface area AI, and maximum R = 0.471 between the rostral middle frontal thickness AI and superior frontal thickness AI.

#### Cohen’s *d* effect sizes

The *t*-statistic for the factor ‘diagnosis’ in each linear mixed effects model was used to calculate Cohen’s *d*, with

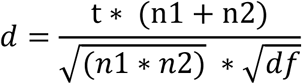

where *n*1 and *n*2 are the number of cases and controls, and *df* the degrees of freedom. The latter was derived from the lme summary table in R, but can also be calculated using *df* = *obs* – (*n*1 + *n*2), where *obs* equals the number of observations, *n*1 the number of groups and *n*2 the number of factors in the model.

The 95% confidence intervals for Cohen’s *d* were calculated using 95% *CI* = *d* ± 1.96 * *SE*, with the standard error (SE) around Cohen’s *d* calculated according to:

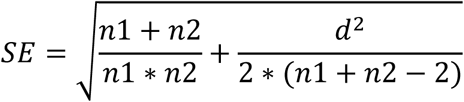

For visualization of cerebral cortical results, Cohen’s d values were loaded into Matlab (version R2016a), and 3D images of left hemisphere inflated cortical and subcortical structures were obtained using FreeSurfer-derived ply files.

#### Power analyses

As each linear model included multiple predictor variables, the power to detect an effect of diagnosis on AI could not be computed exactly, but we obtained an indication of the effect size that would be needed to provide 80% power, had we been using simple t-tests and Bonferroni correction for multiple testing, using the ‘pwr’ command in R. For this purpose, a significance level of 0.0014 (i.e. 0.05/35) was set in the context of multiple testing over the regional and total cortical surface area AIs (N = 35) or thickness AIs (N = 35), and 0.00625 (i.e., 0.05/8) for seven subcortical volume plus lateral ventricle AIs (N = 8). This showed that a difference of roughly Cohen’s *d* = 0.13 would be detectable with 80% power in the cortical analyses, and Cohen’s *d* = 0.12 in the subcortical analyses.

#### Directions of asymmetry changes

For any AIs showing significant effects of diagnosis in the main analysis, linear mixed effects modelling was also performed on the corresponding L and R measures separately, to understand the unilateral changes involved. The models included the same terms as were used in the main analysis of AIs (i.e., *diagnosis, age* and *sex* as fixed factors, and *dataset* as random factor). Again, the Cohen’s *d* effect sizes for diagnosis were calculated based on the *t*-statistics. The raw mean AI values were calculated separately in controls and cases, to describe the reference direction of healthy asymmetry in controls, and whether cases showed reduced, increased, or reversed asymmetry relative to controls.

#### Age- or sex-specific effects

For any AIs having significant effects of diagnosis in the primary analysis, *post hoc* analyses were performed including *age* * *diagnosis* and *sex* * *diagnosis* interaction terms, in separate models. For this purpose, age was coded as a binary factor variable (children < 18y and adults ≥ 18 y) (i.e., using the same criteria as van Rooij *et al.* 2018). Thus the models were as follows: *AI* = *diagnosis* + *factor*(*age*) + *sex* + *factor*(*age*) * *diag* + *random* (= *dataset*), and *AI* = *diagnosis* + *age* + *sex* + *sex* * *diag* + *random* (=*dataset*).

Where we identified an age:diagnosis interaction with *P* < 0.05 (uncorrected), we separated the data for the relevant AI into two subsets by age (i.e. children, adults), and tested the following model within each subset separately: *AI* = *diagnosis* + *sex* + *random* (= *dataset*), to better understand the relation of diagnosis to AI. Likewise, if a sex:diagnosis interaction was found, then a model was fitted separately within each sex, such that *AI* = *diagnosis* + *age* + *random* (= *dataset*).

#### Case-only analysis of IQ

For each AI showing a significant effect of diagnosis in the main analysis, a within-case-only analysis was performed, whereby IQ (as a continuous variable) was considered as a predictor variable for the AI, so that *AI* = *IQ* + *age* + *sex* + *random* (= *dataset*). This was done to understand whether asymmetry changes were a feature of high or low IQ, or occurred regardless of IQ.

#### Autism Diagnostic Observation Schedule (ADOS) severity score

Likewise, for each AI showing a significant effect of diagnosis in the main analysis, a within-case-only analysis was performed incorporating symptom severity based on ADOS score as a predictor variable for AI: *AI* = *ADOS* + *age* + *sex* + *random* (= *dataset*). This was to understand whether the observed asymmetry changes in cases were dependent on ASD severity.

ADOS scores were first normalized using the R package ‘bestNormalize’, which selected the optimal transformation method, based on the lowest Pearson P test statistic for normality, among the Yeo-Johnson, exp(x), log10, square-root, arcsinh and orderNorm transformations (https://github.com/petersonR/bestNormalize). The orderNorm transformation was selected, which performs ordered quantile normalizing (i.e. ranked normalization) using the following formula:

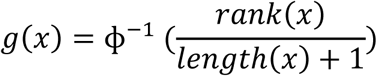

- where Φ refers to the standard normal cumulative distribution function, *rank*(*x*) to the rank of each observation, and *length*(*x*) to the number of observations.

## Results

### Main analyses

#### Significant associations of ASD with brain asymmetry

Out of a total of 78 structural AIs that were investigated (**Tables S3-S5**), 11 showed a significant effect of diagnosis which survived multiple testing correction (**Table 1**). Among these were the total hemispheric average thickness AI (β =-0.0007, *t* = −2.6, *P* = 0.0082) (**Table 1, Table S3**), as well as eight regional cortical thickness AIs, including frontal regions (superior frontal, rostral middle frontal, medial orbitofrontal), temporal regions (fusiform, inferior temporal, superior temporal), and cingulate regions (rostral anterior, isthmus cingulate). One cortical regional surface area AI, namely of the medial orbitofrontal cortex, was significantly associated with diagnosis (β = 0.008, *t* = 4.2, *P* = 0.00003) (**Table 1, Table S4**), and one subcortical volume AI, namely that of the putamen (β = 0.0036, t = 3.6, *P* = 0.00028) (**Table 1, Table S5**).

**Table 1.** Linear mixed model results for regional AIs that survived multiple comparisons correction.

| AI Region | N cases/<br>controls | t-value |  |  | p-value <sup>a</sup> |  |  | Cohen's *d*<br>(95% CI) |
| --- | --- | --- | --- | --- | --- | --- | --- | --- |
|  |  | diag | age | sex | diag | age | sex |  |
| Fusiform thickness | 1605/1618 | 3.98 | -0.20 | 1.37 | 0.0002 | 0.839 | 0.170 | 0.141 (0.07,0.21) |
| Inferiortemporal thickness | 1589/1624 | 3.20 | -1.12 | 0.41 | 0.001 | 0.262 | 0.678 | 0.114 (0.04,0.18) |
| Isthmuscingulate thickness | 1594/1635 | -3.49 | 2.07 | 0.50 | 0.0004 | 0.039 | 0.618 | -0.124 (-0.19,-0.05) |
| Medialorbitofrontal thickness | 1626/1629 | -3.83 | 3.96 | -0.47 | 0.0002 | 8.5×10<sup>-5</sup> | 0.642 | -0.135 (-0.2,-0.07) |
| Rostralanteriorcingulate thickness | 1597/1631 | -3.93 | -0.51 | 0.95 | 9.4×10<sup>-5</sup> | 0.613 | 0.344 | -0.14 (-0.21,-0.07) |
| Rostralmiddlefrontal thickness | 1597/1642 | -3.99 | -1.73 | 0.98 | 0.0001 | 0.084 | 0.328 | -0.142 (-0.21,-0.07) |
| Superiorfrontal thickness | 1604/1626 | -4.52 | 0.46 | -0.65 | 4.5×10<sup>-6</sup> | 0.642 | 0.513 | -0.16 (-0.23,-0.09) |
| Superiortemporal thickness | 1573/1644 | -2.54 | 0.47 | 1.47 | 0.011 | 0.637 | 0.141 | -0.09 (-0.16,-0.02) |
| Total average thickness | 1604/1645 | -2.65 | -0.51 | 0.85 | 0.008 | 0.613 | 0.396 | -0.094 (-0.16,-0.02) |
| Medialorbitofrontal surface area | 1608/1640 | 4.18 | 1.03 | -1.13 | 3.4×10<sup>-5</sup> | 0.305 | 0.257 | 0.148 (0.08,0.22) |
| Putamen volume | 1559/1652 | 3.64 | 3.05 | 1.03 | 0.0004 | 0.002 | 0.304 | 0.13 (0.06,0.2) |

<sup>a</sup> Unadjusted P values are shown. Note that this table only includes AIs for which the effect of diagnosis was significant after FDR correction. Results for all AIs are in supplementary information.

Nominally significant effects of diagnosis on AIs (i.e., with *P* < 0.05, but which did not survive multiple comparison correction), were observed for the inferior parietal thickness AI (β = −0.0017, *t* = −2.4, *P* = 0.015) (**Table S3**), the fusiform surface area AI (β = −0.0040, *t* = − 2.4, *P* = 0.016), precentral surface area AI (β = −0.0031, *t* = −2.5, *P* = 0.011), lateral orbitofrontal surface area AI (β = −0.0035, *t* = −2.5, *P* = 0.012), and the *pars orbitalis* surface area AI (β = 0.0044, *t* = 2.2, *P* = 0.025)(**Table S4**),

When we repeated the analysis without excluding outliers, 7 out of the 11 previously significant AIs remained significant after FDR correction (AIs of the total average thickness, isthmus cingulate thickness, superior temporal thickness and medial orbitofrontal surface area did not survive multiple testing correction (**Table S6**)). When we repeated the analysis including a non-linear effect for age, all of the 11 AIs that had shown significant effects of diagnosis in the main analysis remained significant (**Table S6**). Finally, when we excluded the two youngest datasets (i.e., FSM and MYAD) that may have been more difficult for FreeSurfer to segment, all AIs that had shown significant effects of diagnosis in the main analysis remained significant, except for the superior temporal thickness AI (**Table S6**).

#### Magnitudes and directions of asymmetry changes

Cohen’s *d* effect sizes of the associations between AIs and diagnosis are visualized per region in Figure 1, as derived from the main analysis. Effect sizes were small, ranging from −0.16 (superior frontal thickness AI) to 0.15 (medial orbitofrontal surface area AI) (**Table 1**, **Tables S3-S5**).

**Figure 1.**
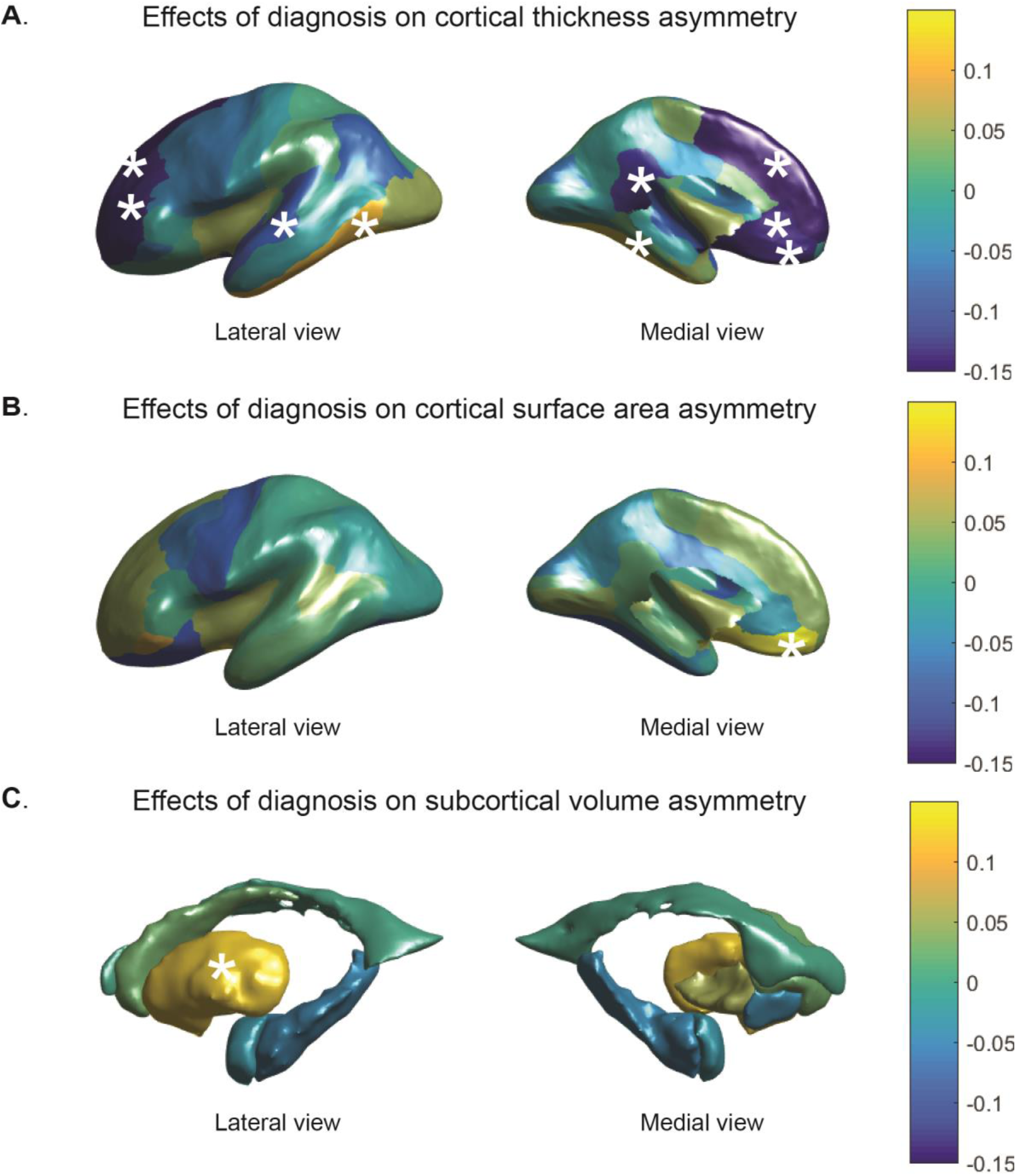
Cohen’s *d* effect sizes of the associations between diagnosis and AIs, for (**A**) regional cortical thickness measures (**B**) cortical surface areas, and (**C**) subcortical volumes. Values are overlaid on left hemisphere inflated brains. Positive Cohen’s *d* values (*yellow*) indicate mean shifts towards greater leftward or reduced rightward asymmetry in cases, and negative Cohen’s *d* values (*blue*) indicate mean shifts towards greater rightward asymmetry or reduced leftward asymmetry in cases. Regions that showed a significant association between diagnosis and AI after multiple testing correction, are indicated with stars.

The raw mean AIs in ASD individuals and controls are listed in **Table S7**. There was reduced leftward asymmetry in total hemispheric average cortical thickness (*d* = −0.094) (**Table 1, Figure 1**) in ASD compared to controls, driven by a larger decrease in left (*d*=-0.148) than right (*d*=-0.111) thickness (**Table S7**) in ASD. The difference in superior temporal thickness asymmetry was also driven by a larger decrease on the left (*d* = −0.130) than on the right (*d* = −0.065) in cases (**Table S7**), resulting in an increased rightward AI in ASD (*d* = −0.090) (**Table 1, Figure 1A**).

Four regions showed reduced left-side thickness together with increased right-side thickness in ASD, leading to decreased leftward asymmetry. These were the isthmus cingulate (AI *d* = − 0.12, left *d* = −0.027, right *d*=0.091), rostral anterior cingulate (AI *d* = −0.140, left d = −0.128, right d= 0.018), rostral middle frontal (AI d = −0.142, left *d* = −0.018, right *d* = 0.061), and superior frontal (AI *d* = −0.160, left *d* = −0.069, right 0.018) (**Table 1, Table S7, Figure 1A**).

Two regions showed a smaller reduction in left than right sided thickness in ASD, resulting in reduced rightward asymmetry. These were the fusiform (AI *d* = 0.141, left *d* = −0.201, right d= −0.285) and inferior temporal cortex (AI *d* = 0.114, left *d* = −0.143, right *d*= −0.218) (**Table 1, Table S7, Figure 1A**).

The thickness of the right medial orbitofrontal cortex was increased in ASD, while being virtually unchanged on the left, resulting in a reversal in thickness asymmetry from leftward in controls to rightward in ASD (AI *d* = −0.135, left *d* = 0.014, right *d* = 0.121) (**Table 1, Table S7, Figure 1A**). Similarly, surface area asymmetry of this region was significantly associated with diagnosis, whereby the ASD group had reduced rightward asymmetry, driven by an increase in left area and a decrease in right area (AI *d* = 0.15, left *d* = 0.094, right *d* = − 0.019) (**Table 1, Table S7, Figure 1B**).

Notably, all significant changes in cortical thickness asymmetry, with the exception of the superior temporal cortex, involved a decrease in the magnitude of asymmetry in ASD compared to controls (**Table S7**), whether it was reduced leftward, reduced rightward, or reversed asymmetry.

For the putamen, cases showed an increased leftward volume asymmetry compared to controls (*d*=0.130) (**Table 1, Figure 1C**), driven by a larger decrease in right volume (*d* = − 0.084) than left volume (d = −0.022) in ASD (**Table S7**).

#### Age- and sex-limited effects

The distributions of age and sex across all datasets are plotted in Figure S1A,B. None of the 11 AIs with significant case-control differences in the main analysis were associated with a significant age by diagnosis interaction effect (**Table S8**). A putative sex by diagnosis interaction was observed for the rostral anterior cingulate thickness AI (p_sex*diag_ = 0.00114) (**Table S8**). The AI for this regional thickness was associated with diagnosis in males (p = 9 × 10^−7^) but not females (p = 0.22) (**Table S8**).

#### Analysis of IQ within cases

The distribution of IQ within the ASD group is shown in Figure S1C. Out of the 11 AIs which showed significant case-control differences in the main analysis, only one showed an association with IQ (uncorrected P < 0.05), which was the rostral anterior cingulate thickness AI (β = 0.00024, t = 3.3, p = 0.0009) (**Table S9**). The positive direction of this effect indicates that primarily cases with lower IQ show reduced leftward asymmetry of this regional thickness.

#### ADOS severity scores

The distribution of ADOS severity scores is plotted in Figure S1D.

Out of the 11 AIs showing significant case-control differences in the main analysis, only the isthmus cingulate thickness AI showed a significant association with the ADOS severity score (β =0.0041, t = 2.6, p = 0.011) (**Table S2**). The positive direction of the effect suggests that primarily cases with low ASD severity have reduced leftward asymmetry of this regional thickness.

## Discussion

In this, the largest study to date of brain asymmetry in ASD, we mapped differences in brain asymmetry between participants with ASD and typically developing controls, in a collection of 54 international datasets via the ENIGMA Consortium. We had 80% statistical power to detect Cohen’s *d* effect sizes in the range of 0.12 to 0.13. We found an overall decrease in leftward asymmetry of total average cortical thickness, that is to say, considered at a hemispheric level over all cortical regions in autistic individuals. Several specific regional thickness asymmetries - located broadly over the cortex - were also significantly altered in ASD. These included frontal, temporal and cingulate regions. The magnitude of all regional thickness asymmetries, with the exception of the superior temporal cortex, was decreased in ASD compared to controls, whether it was reduced leftward, reduced rightward, or reversed average asymmetry. Rightward asymmetry of the medial orbitofrontal surface area was also decreased in cases. In addition, cases showed an increase in leftward asymmetry of putamen volume, compared to controls.

Previous MRI studies of cerebral cortical asymmetries in ASD, based on much smaller datasets, and using diverse methods for image analysis, suggested variable differences (23, 24), or no differences (21, 25). Our findings partly support a previously reported, generalized reduction of leftward asymmetry (23), as most of the significantly altered regional asymmetries involved right-side thickness increases accompanied by left-side decreases, or else larger left-side than right-side decreases. However, two of the nine significantly altered regional thickness asymmetries in our analysis involved shifts leftwards in ASD, driven by more prominent right-than left-side decreases, i.e., the fusiform and inferior temporal cortex. Thus the directional change of thickness asymmetry can depend on the specific region, albeit that the overall magnitude of thickness asymmetry is most likely to be reduced in ASD.

The significant associations of diagnosis with asymmetry in the present study were all weak (Cohen’s *d* −0.16 to 0.15), indicating that altered structural brain asymmetry is unlikely to be a useful predictor for ASD. Prior studies using smaller samples were underpowered in this context. However, the effect sizes were comparable to those reported by recent, large-scale studies of bilateral disorder-related changes in brain structure, in which asymmetry was not studied, including for ASD (8) as well as ADHD (32), schizophrenia (31), OCD (26, 27), posttraumatic stress disorder (28), and major depressive disorder (29, 30). It has become increasingly clear that anatomical differences between ASD and control groups are very small relative to the large within-group variability that is observed (43).

Our findings may inform understanding of the neurobiology of ASD. Multi-regional reduction of cortical thickness asymmetry in ASD fits with the concept that laterality is an important organizing feature of the healthy human brain for multiple aspects of complex cognition, and is susceptible to disruption in disorders (e.g. (10, 44)). Left-right asymmetry facilitates the development of localized and specialized modules in the brain, which can then have dominant control of behaviour (45, 46). Notably, many of the cortical regions highlighted here are involved in diverse social cognitive processes, including sensory processing (fusiform and superior temporal gyri), cognitive and emotional control (anterior cingulate) and reward evaluation (orbitofrontal cortex, ventral striatum) (47). However, the roles of these brain structures are by no means restricted to social behaviour. As we found altered asymmetry of various additional regions, our findings suggest broader disruption of lateralized neurodevelopment as part of the ASD phenotype.

The medial orbitofrontal cortex was the only region that showed significantly altered asymmetry of both thickness and surface area in ASD, suggesting that disrupted laterality of this region might be particularly important in ASD. The orbitofrontal cortex may be involved in repetitive and stereotyped behaviours in ASD, due to its roles in executive functions (48). Prior studies have reported lower cortical thickness in the left medial orbitofrontal gyrus in ASD (49), altered patterning of gyri and sulci in the right orbitofrontal cortex (50), and altered asymmetry in frontal regions globally (19, 25). These studies were in much smaller sample sizes than used here.

As regards the fusiform cortex, a previous study reported an association between higher ASD symptom severity and increased rightward grey matter volume asymmetry (24). We found lower rather than higher rightward thickness asymmetry in ASD in this region The fusiform gyrus is involved in facial memory among other functions, which is important for social interactions (51).

The altered volume asymmetry of the putamen in cases may be related to its role in repetitive and restricted behaviours in ASD. One study reported that differences in striatal growth trajectories were correlated with circumscribed interests and insistence on sameness (52). The striatum is connected with lateral and orbitofrontal regions of the cortex via lateral-frontal-striatal reward circuitry, and this circuitry might be dysfunctional in ASD (53). For example, boys with ASD (and also boys with OCD), exhibited decreased activation of the ventral striatum and lateral inferior/orbitofrontal cortex during outcome anticipation compared with typically developing controls (53).

Although the reasons for asymmetrical alterations in many of the structures implicated here are unclear, our findings suggest altered neurodevelopment affecting these structures in ASD. Further research is necessary to clarify the functional relevance and relationships between altered asymmetry and autism spectrum disorder. The findings we report in this large-scale study did sometimes not concur with prior, smaller studies. This may be due to limited statistical power in the earlier studies, which may have resulted in false positive findings.

However, the cortical atlas that we used did not have perfect equivalents for regions defined in many of the earlier studies, and we did not consider gyral/sulcal patterns, or grey matter volumes as such. Rather, we studied regional cortical thickness and surface areas as distinct measures, which together drive grey matter volumetric measures, but have been shown to vary relatively independently (54). As such, separate analyses are well motivated.

Furthermore, discrepancies with earlier studies may be related to age differences, and differences in clinical features of the disorder arising from case recruitment and diagnosis. We included subjects from the entire ASD severity spectrum, with a broad range of ages, IQs, and of both sexes.

Interestingly, *post hoc* analysis of asymmetries that had shown significant case-control effects in the main analysis, showed that the effects were not influenced by age (**Table S8**). Thus altered asymmetry in ASD appears to have an early neurodevelopmental onset and then remain stable. Only one effect of diagnosis on regional asymmetry was influenced by sex, i.e. the rostral anterior cingulate thickness asymmetry, which was altered in males but not females. This same regional asymmetry was primarily altered in lower versus higher IQ cases. This may therefore be an alteration of cortical asymmetry that is relatively specific to an ASD subgroup, i.e., lower-performing males. As regards symptom severity, thickness asymmetry of the isthmus of the cingulate was associated with the ADOS score, such that the lower severity cases tended to have the most altered asymmetry. However, these post hoc findings remain tentative in the context of multiple testing, and are reported here for descriptive purposes only.

There were additional data available on handedness, medication use and comorbidities for some datasets, but not all. We did not investigate possible associations of these variables with asymmetries, due to the reduced sample size and increased multiple testing. As mentioned above, data on comorbidities were only available for 54 of the ASD subjects, precluding a high-powered analysis of this issue. Handedness had no significant effect on the same brain asymmetry measures as analyzed here, in a separate study of healthy individuals comprising more than 15,000 participants (55, 56). The previous ENIGMA study from the ASD working group did not detect effects of medication use on bilateral brain changes in ASD (8).

In contrast to some prior studies of ASD, we did not adjust for IQ as a covariate effect in our main, case-control analysis. Rather we carried out post hoc analysis, within cases only, of possible associations between IQ and brain asymmetries. This was because lower average IQ was clearly part of the ASD phenotype in our total combined dataset (**Figure S1C**), so that including IQ as a confounding factor in case-control analysis might have reduced the power to detect an association of diagnosis with asymmetry. This would occur if underlying susceptibility factors contribute both to altered asymmetry and reduced IQ, as part of the ASD phenotype.

The Desikan atlas (38) was derived from manual segmentations of sets of reference brain images. Accordingly, the mean regional asymmetries in our samples partly reflect left-right differences present in the reference dataset used to construct the atlas. For detecting cerebral asymmetries with automated methods, some groups have chosen to work from artificially created, left-right symmetrical atlases, e.g. (57). However, our study was focused on comparing *relative* asymmetry between groups. The use of a ‘real-world’ asymmetrical atlas had the advantage that regional identification was likely to be more accurate for structures that are asymmetrical both in the atlas and, on average, in our datasets. By defining the regions of interest in each hemisphere based on each hemisphere’s own particular features, such as its sulcal and gyral geometry, we could then obtain the corresponding relationships between hemispheres. To this end, we used data from the automated labeling program within FreeSurfer for subdividing the human cerebral cortex. The labeling system incorporates hemisphere-specific information on sulcal and gyral geometry with spatial information regarding the locations of brain structures, and shows a high accuracy when compared to manual labeling results (38). Thus, reliable measures of each region can be extracted for each subject, and regional asymmetries then accurately assessed.

Although a single image analysis pipeline was applied to all datasets, heterogeneity of imaging protocols was a limitation of this study. There were substantial differences between datasets in the average asymmetry measured for some regions, which may be due in part to different scanner characteristics, as well as differences in patient profiles. Although we corrected for dataset as a random effect, the between-centre variability may have resulted in reduced statistical power relative to an equally sized single-centre study. However, no single centre has been able to collect such large samples alone. In addition, multi-centre studies may better represent real-world heterogeneity, with more generalizable findings than single-centre studies (58)

The cross-sectional design limits our capacity to make causal inferences between diagnosis and asymmetry. ASD is highly heritable, with meta-analytic heritability estimates ranging from 64% to 91% (59). Likewise, some of the brain asymmetry measures examined here have heritabilities as high as roughly 25% (55, 56). Future studies are required to investigate shared genetic contributions to ASD and variation in brain structural asymmetry. These could help to disentangle cause-effect relations between ASD and brain structural asymmetry. Given the high comorbidity of ASD with other disorders, such as ADHD, OCD, and schizophrenia (60), cross-disorder analyses incorporating between-disorder genetic correlations may be informative.

In conclusion, large-scale analysis of brain asymmetry in ASD revealed primarily cortical thickness effects, which were significant but small. Our study illustrates how high-powered and systematic studies can yield much needed clarity in human clinical neuroscience, where prior smaller and methodologically diverse studies were inconclusive.

## Acknowledgements

We thank the participants of all studies who have contributed data to the ENIGMA-ASD working group (http://enigma.ini.usc.edu/ongoing/enigma-asd-working-group/) (8). This research was funded by the Max Planck Society (Germany). This study was further supported by the ENIGMA Center for Worldwide Medicine, Imaging & Genomics grant (NIH U54 EB020403) to Paul Thompson, and further supported by the Innovative Medicines Initiative Joint Undertaking under grant agreement number 115300 (EU-AIMS) and 777394 (AIMS-2-TRIALS), resources of which are composed of financial contribution from the European Union’s Seventh Framework Programme and Horizon2020 programmes and the European Federation of Pharmaceutical Industries and Associations (EFPIA) companies’ in kind contribution. The Canadian samples were collected as part of the POND network funded by the Ontario Brain Institute (grant IDS-I l-02 to Anagnostou / Lerch).

## Disclosures

Dr. Anagnostou has served as a consultant or advisory board member for Roche and Takeda; she has received funding from the Alva Foundation, Autism Speaks, Brain Canada, the Canadian Institutes of Health Research, the Department of Defense, the National Centers of Excellence, NIH, the Ontario Brain Institute, the Physicians’ Services Incorporated (PSI) Foundation, Sanofi-Aventis, and SynapDx, as well as in-kind research support from AMO Pharma; she receives royalties from American Psychiatric Press and Springer and an editorial honorarium from Wiley. Her contribution is on behalf of the POND network. Dr. Arango has served as a consultant for or received honoraria or grants from Acadia, Abbott, Amgen, CIBERSAM, Fundación Alicia Koplowitz, Instituto de Salud Carlos III, Janssen-Cilag, Lundbeck, Merck, Instituto de Salud Carlos III (co-financed by the European Regional Development Fund “A way of making Europe,” CIBERSAM, the Madrid Regional Government [S2010/BMD-2422 AGES], the European Union Structural Funds, and the European Union Seventh Framework Programmeunder grant agreements FP7-HEALTH-2009-2.2.1-2-241909, FP7-HEALTH-2009-2.2.1-3-242114, FP7-HEALTH-2013-2.2.1-2-603196, and FP7-HEALTH-2013-2.2.1-2-602478), Otsuka, Pfizer, Roche, Servier, Shire, Takeda, and Schering-Plough. Dr. Freitag has served as a consultant for Desitin regarding issues on ASD. Dr. Di Martino is a coauthor of the Italian version of the Social Responsiveness Scale, for which she may receive royalties. Her contribution is on behalf of the ABIDE and ABIDE-II consortia. Dr. Rubia has received speaking honoraria fromEli Lilly, Medice, and Shire, and a grant from Shire for another project. Dr. Buitelaar has served as a consultant, advisory board member, or speaker for Eli Lilly, Janssen-Cilag, Lundbeck, Medice, Novartis, Servier, Shire, and Roche, and he has received research support fromRoche and Vifor. The other authors report no financial relationships with commercial interests.

## References

1. American Psychiatric Association (2013): Diagnostic and statistical manual of mental disorders (5th ed.). Washington, DC.

2. Xu G, Strathearn L, Liu B, O’Brien M, Kopelman TG, Zhu J, et al. (2018): Prevalence and Treatment Patterns of Autism Spectrum Disorder in the United States, 2016. JAMA pediatrics.

3. Loth E, Murphy DG, Spooren W (2016): Defining Precision Medicine Approaches to Autism Spectrum Disorders: Concepts and Challenges. Front Psychiatry. 7:188.

4. Li D, Karnath HO, Xu X (2017): Candidate Biomarkers in Children with Autism Spectrum Disorder: A Review of MRI Studies. Neuroscience bulletin. 33:219–237.

5. Rommelse N, Buitelaar JK, Hartman CA (2017): Structural brain imaging correlates of ASD and ADHD across the lifespan: a hypothesis-generating review on developmental ASD-ADHD subtypes. Journal of neural transmission (Vienna, Austria: 1996). 124:259–271.

6. Biberacher V, Schmidt P, Keshavan A, Boucard CC, Righart R, Samann P, et al. (2016): Intra- and interscanner variability of magnetic resonance imaging based volumetry in multiple sclerosis. Neuroimage. 142:188–197.

7. Jeste SS, Geschwind DH (2014): Disentangling the heterogeneity of autism spectrum disorder through genetic findings. Nature reviews Neurology. 10:74–81.

8. van Rooij D, Anagnostou E, Arango C, Auzias G, Behrmann M, Busatto GF, et al. (2018): Cortical and Subcortical Brain Morphometry Differences Between Patients With Autism Spectrum Disorder and Healthy Individuals Across the Lifespan: Results From the ENIGMA ASD Working Group. Am J Psychiatry. 175:359–369.

9. Duboc V, Dufourcq P, Blader P, Roussigne M (2015): Asymmetry of the Brain: Development and Implications. Annu Rev Genet. 49:647–672.

10. Renteria ME (2012): Cerebral asymmetry: a quantitative, multifactorial, and plastic brain phenotype. Twin Res Hum Genet. 15:401–413.

11. Toga AW, Thompson PM (2003): Mapping brain asymmetry. Nat Rev Neurosci. 4:37–48.

12. Knaus TA, Silver AM, Kennedy M, Lindgren KA, Dominick KC, Siegel J, et al. (2010): Language laterality in autism spectrum disorder and typical controls: a functional, volumetric, and diffusion tensor MRI study. Brain and language. 112:113–120.

13. Kleinhans NM, Muller RA, Cohen DN, Courchesne E (2008): Atypical functional lateralization of language in autism spectrum disorders. Brain research. 1221:115–125.

14. Lindell AK, Hudry K (2013): Atypicalities in cortical structure, handedness, and functional lateralization for language in autism spectrum disorders. Neuropsychol Rev. 23:257–270.

15. Cardinale RC, Shih P, Fishman I, Ford LM, Muller RA (2013): Pervasive rightward asymmetry shifts of functional networks in autism spectrum disorder. JAMA Psychiatry. 70:975–982.

16. Markou P, Ahtam B, Papadatou-Pastou M (2017): Elevated Levels of Atypical Handedness in Autism: Meta-Analyses. Neuropsychol Rev. 27:258–283.

17. Rysstad AL, Pedersen AV (2018): There Are Indeed More Left-Handers Within the Autism Spectrum Disorder Compared with in the General Population, but the Many Mixed-Handers Is the More Interesting Finding. J Autism Dev Disord.

18. Gabard-Durnam L, Tierney AL, Vogel-Farley V, Tager-Flusberg H, Nelson CA (2015): Alpha asymmetry in infants at risk for autism spectrum disorders. J Autism Dev Disord. 45:473–480.

19. Conti E, Calderoni S, Gaglianese A, Pannek K, Mazzotti S, Rose S, et al. (2016): Lateralization of Brain Networks and Clinical Severity in Toddlers with Autism Spectrum Disorder: A HARDI Diffusion MRI Study. Autism research: official journal of the International Society for Autism Research. 9:382–392.

20. Carper RA, Treiber JM, DeJesus SY, Muller RA (2016): Reduced Hemispheric Asymmetry of White Matter Microstructure in Autism Spectrum Disorder. Journal of the American Academy of Child and Adolescent Psychiatry. 55:1073–1080.

21. Joseph RM, Fricker Z, Fenoglio A, Lindgren KA, Knaus TA, Tager-Flusberg H (2014): Structural asymmetries of language-related gray and white matter and their relationship to language function in young children with ASD. Brain Imaging Behav. 8:60–72.

22. Wei L, Zhong S, Nie S, Gong G (2018): Aberrant development of the asymmetry between hemispheric brain white matter networks in autism spectrum disorder. Eur Neuropsychopharmacol. 28:48–62.

23. Floris DL, Lai MC, Auer T, Lombardo MV, Ecker C, Chakrabarti B, et al. (2016): Atypically rightward cerebral asymmetry in male adults with autism stratifies individuals with and without language delay. Hum Brain Mapp. 37:230–253.

24. Dougherty CC, Evans DW, Katuwal GJ, Michael AM (2016): Asymmetry of fusiform structure in autism spectrum disorder: trajectory and association with symptom severity. Molecular autism. 7:28.

25. Knaus TA, Tager-Flusberg H, Mock J, Dauterive R, Foundas AL (2012): Prefrontal and occipital asymmetry and volume in boys with autism spectrum disorder. Cognitive and behavioral neurology: official journal of the Society for Behavioral and Cognitive Neurology. 25:186–194.

26. Boedhoe, Schmaal L, Abe Y, Ameis SH, Arnold PD, Batistuzzo MC, et al. (2017): Distinct Subcortical Volume Alterations in Pediatric and Adult OCD: A Worldwide Meta- and Mega-Analysis. Am J Psychiatry. 174:60–69.

27. Boedhoe, Schmaal L, Abe Y, Alonso P, Ameis SH, Anticevic A, et al. (2018): Cortical Abnormalities Associated With Pediatric and Adult Obsessive-Compulsive Disorder: Findings From the ENIGMA Obsessive-Compulsive Disorder Working Group. Am J Psychiatry. 175:453–462.

28. Logue MW, van Rooij SJH, Dennis EL, Davis SL, Hayes JP, Stevens JS, et al. (2018): Smaller Hippocampal Volume in Posttraumatic Stress Disorder: A Multisite ENIGMA-PGC Study: Subcortical Volumetry Results From Posttraumatic Stress Disorder Consortia. Biol Psychiatry. 83:244–253.

29. Schmaal L, Hibar DP, Samann PG, Hall GB, Baune BT, Jahanshad N, et al. (2017): Cortical abnormalities in adults and adolescents with major depression based on brain scans from 20 cohorts worldwide in the ENIGMA Major Depressive Disorder Working Group. Mol Psychiatry. 22:900–909.

30. Schmaal L, Veltman DJ, van Erp TG, Samann PG, Frodl T, Jahanshad N, et al. (2016): Subcortical brain alterations in major depressive disorder: findings from the ENIGMA Major Depressive Disorder working group. Mol Psychiatry. 21:806–812.

31. van Erp TGM, Walton E, Hibar DP, Schmaal L, Jiang W, Glahn DC, et al. (2018): Cortical Brain Abnormalities in 4474 Individuals With Schizophrenia and 5098 Control Subjects via the Enhancing Neuro Imaging Genetics Through Meta Analysis (ENIGMA) Consortium. Biol Psychiatry. 84:644–654.

32. Hoogman M, Bralten J, Hibar DP, Mennes M, Zwiers MP, Schweren LS, et al. (2017): Subcortical brain volume differences in participants with attention deficit hyperactivity disorder in children and adults: a cross-sectional mega-analysis. The lancet Psychiatry. 4:310–319.

33. Munafo MR, Flint J (2010): How reliable are scientific studies? Br J Psychiatry. 197:257–258.

34. Button KS, Ioannidis JP, Mokrysz C, Nosek BA, Flint J, Robinson ES, et al. (2013): Power failure: why small sample size undermines the reliability of neuroscience. Nat Rev Neurosci. 14:365– 376.

35. Kurth F, Gaser C, Luders E (2015): A 12-step user guide for analyzing voxel-wise gray matter asymmetries in statistical parametric mapping (SPM). Nature protocols. 10:293–304.

36. Leroy F, Cai Q, Bogart SL, Dubois J, Coulon O, Monzalvo K, et al. (2015): New human-specific brain landmark: the depth asymmetry of superior temporal sulcus. Proc Natl Acad Sci U S A. 112:1208–1213.

37. Lord C, Risi S, Lambrecht L, Cook EH, Jr., Leventhal BL, DiLavore PC, et al. (2000): The autism diagnostic observation schedule-generic: a standard measure of social and communication deficits associated with the spectrum of autism. J Autism Dev Disord. 30:205–223.

38. Desikan RS, Segonne F, Fischl B, Quinn BT, Dickerson BC, Blacker D, et al. (2006): An automated labeling system for subdividing the human cerebral cortex on MRI scans into gyral based regions of interest. Neuroimage. 31:968–980.

39. Fischl B (2012): FreeSurfer. Neuroimage. 62:774–781.

40. Pinheiro J BD, DebRoy S, Sarkar D and R Core Team (2018). nlme: Linear and Nonlinear Mixed Effects Models. R package version 3.1–137, https://CRAN.R-project.org/package=nlme.

41. Chambers JM, Hastie TJ (1992): Statistical models in S. Pacific Grove, California, USA, Wadsworth & Brooks/Cole.

42. Benjamini Y, Hochberg Y (1995): Controlling the False Discovery Rate - A Practical and Powerful Approach to Multiple Testing. J R Stat Soc Ser B-Methodol. 57:289–300.

43. Haar S, Berman S, Behrmann M, Dinstein I (2016): Anatomical Abnormalities in Autism? Cereb Cortex. 26:1440–1452.

44. Francks C (2015): Exploring human brain lateralization with molecular genetics and genomics. Ann N Y Acad Sci. 1359:1–13.

45. Geschwind N, Galaburda AM (1985): Cerebral lateralization: Biological mechanisms, associations, and pathology: i. a hypothesis and a program for research. Archives of Neurology. 42:428–459.

46. Rogers LJ, Zucca P, Vallortigara G (2004): Advantages of having a lateralized brain. Proc Biol Sci. 271 Suppl 6:S420–422.

47. Adolphs R (2009): The social brain: neural basis of social knowledge. Annual review of psychology. 60:693–716.

48. Ecker C, Bookheimer SY, Murphy DG (2015): Neuroimaging in autism spectrum disorder: brain structure and function across the lifespan. The Lancet Neurology. 14:1121–1134.

49. Jiao Y, Chen R, Ke X, Chu K, Lu Z, Herskovits EH (2010): Predictive models of autism spectrum disorder based on brain regional cortical thickness. Neuroimage. 50:589–599.

50. Watanabe H, Nakamura M, Ohno T, Itahashi T, Tanaka E, Ohta H, et al. (2014): Altered orbitofrontal sulcogyral patterns in adult males with high-functioning autism spectrum disorders. Social cognitive and affective neuroscience. 9:520–528.

51. Trontel HG, Duffield TC, Bigler ED, Froehlich A, Prigge MB, Nielsen JA, et al. (2013): Fusiform correlates of facial memory in autism. Behavioral sciences (Basel, Switzerland). 3:348–371.

52. Langen M, Bos D, Noordermeer SD, Nederveen H, van Engeland H, Durston S (2014): Changes in the development of striatum are involved in repetitive behavior in autism. Biol Psychiatry. 76:405–411.

53. Carlisi CO, Norman L, Murphy CM, Christakou A, Chantiluke K, Giampietro V, et al. (2017): Shared and Disorder-Specific Neurocomputational Mechanisms of Decision-Making in Autism Spectrum Disorder and Obsessive-Compulsive Disorder. Cereb Cortex. 27:5804–5816.

54. Panizzon MS, Fennema-Notestine C, Eyler LT, Jernigan TL, Prom-Wormley E, Neale M, et al. (2009): Distinct genetic influences on cortical surface area and cortical thickness. Cereb Cortex. 19:2728–2735.

55. Guadalupe T, Mathias SR, vanErp TG, Whelan CD, Zwiers MP, Abe Y, et al. (2016): Human subcortical brain asymmetries in 15,847 people worldwide reveal effects of age and sex. Brain Imaging Behav.

56. Kong XZ, Mathias SR, Guadalupe T, Glahn DC, Franke B, Crivello F, et al. (2018): Mapping cortical brain asymmetry in 17,141 healthy individuals worldwide via the ENIGMA Consortium. Proc Natl Acad Sci U S A.

57. Kawasaki Y, Suzuki M, Takahashi T, Nohara S, McGuire PK, Seto H, et al. (2008): Anomalous cerebral asymmetry in patients with schizophrenia demonstrated by voxel-based morphometry. Biol Psychiatry. 63:793–800.

58. Costafreda SG (2009): Pooling FMRI data: meta-analysis, mega-analysis and multi-center studies. Frontiers in neuroinformatics. 3:33.

59. Tick B, Bolton P, Happe F, Rutter M, Rijsdijk F (2016): Heritability of autism spectrum disorders: a meta-analysis of twin studies. Journal of child psychology and psychiatry, and allied disciplines. 57:585–595.

60. Sharma SR, Gonda X, Tarazi FI (2018): Autism Spectrum Disorder: Classification, diagnosis and therapy. Pharmacology & therapeutics.

