## Supplementary for "Altered structural brain asymmetry in autism spectrum disorder: large-scale analysis via the ENIGMA Consortium" (from another version of this preprint: version 1, 2019-03-09)

**Table S1:** Characteristics of the different datasets used in the mega-analysis

| Sample name | N total | N cases (M/F) | N controls (M/F) | Median age (range) | Scanner type | Field strength |
|---|---|---|---|---|---|---|
| ABIDE_CALTECH | 31 | 43478 | 43568 | 23.4 (17.5, 56.2) | Siemens Trio | 3T |
| ABIDE_KKI | 21 | 7/0 | 43535 | 10.57 (8.43, 12.76) | Philips Achieva | 3T |
| ABIDE_LEUVEN_1 | 29 | 14/0 | 15/0 | 22 (18, 32) | Philips Interna | 3T |
| ABIDE_LEUVEN_2 | 35 | 43536 | 43600 | 14.2 (12.1, 16.9) | Philips Interna | 3T |
| ABIDE_MAX_MUN | 57 | 43545 | 43584 | 26 (7, 58) | Siemens Verio | 3T |
| ABIDE_NYU | 186 | 68/11 | 81/26 | 13.61 (6.47, 39.1) | Siemens Allegra | 3T |
| ABIDE_OHSU | 18 | 7/0 | 11/0 | 10.01 (8.2, 12.65) | Siemens Trio | 3T |
| ABIDE_OLIN | 36 | 43541 | 43510 | 17 (10, 24) | Siemens Allegra | 3T |
| ABIDE_PITT | 58 | 43611 | 43578 | 16.99 (9.33, 35.2) | Siemens Allegra | 3T |
| ABIDE_SBL | 30 | 15/0 | 15/0 | 33.5 (20, 64) | Philips Interna | 3T |
| ABIDE_SDSU | 37 | 43479 | 43632 | 14.75 (8.67, 37.7) | GE MR750 | 3T |
| ABIDE_STANFORD | 40 | 43571 | 43571 | 9.418 (7.528, 12.94) | GR Signa | 3T |
| ABIDE_TCD | 55 | 43489 | 30/0 | 15.91 (9.3, 25.91) | Philips Achieva | 3T |
| ABIDE_UM_1 | 130 | 50/14 | 43/23 | 12.3 (8.07, 20.9) | GE Signa | 3T |
| ABIDE_UM_2 | 31 | 43479 | 43480 | 14.8 (11.14, 26.8) | GE Signa | 3T |
| ABIDE_USM | 101 | 59/0 | 42/0 | 19.63 (8.2, 50.22) | Siemens Trio | 3T |
| ABIDE_YALE | 55 | 43697 | 43696 | 12.75 (7, 17.83) | Siemens Magnetom | 3T |
| ABIDEII-BNI | 58 | 29/0 | 29/0 | 43 (18, 64) | Philips Ingenia | 3T |
| ABIDEII-EMC | 54 | 43607 | 43607 | 8.156 (6.212, 10.66) | GE MR750 | 3T |
| ABIDEII-ETH | 37 | 13/0 | 24/0 | 22.33 (13.83, 30.67) | Philips Achieva | 3T |
| ABIDEII-GU | 106 | 43/8 | 28/27 | 10.61 (8.063, 13.91) | Siemens TriTim | 3T |
| ABIDEII-IP | 56 | 43691 | 44896 | 18.4 (6.051, 46.6) | Siemens TriTim | 1.5T |
| ABIDEII-IU | 40 | 43571 | 43600 | 22 (17, 54) | Philips Achieva | 3T |
| ABIDEII-KKI | 211 | 41/15 | 99/56 | 10.3 (8.014, 12.99) | Philips Achieva | 3T |
| ABIDEII-KUL* | 28 | 28/0 | - | 24 (18, 35) | Philips Achieva | 3T |
| ABIDEII-NYU_1 | 78 | 43/5 | 43524 | 8.45 (5.22, 34.76) | Siemens Allegra | 3T |
| ABIDEII-NYU_2* | 27 | 43548 | - | 6.96 (5.128, 8.844) | Siemens Allegra | 3T |
| ABIDEII-OHSU | 93 | 43676 | 27/29 | 11 (7, 15) | Siemens Skyra | 3T |
| ABIDEII-OILH | 59 | 43575 | 20/15 | 23 (18, 31) | Siemens TriTim | 3T |
| ABIDEII-SDSU | 57 | 43672 | 43518 | 13 (7.4, 18) | GE MR750 | 3T |
| ABIDEII-TCD | 43 | 21/0 | 22/0 | 14.75 (10, 20) | Philips Achieva | 3T |
| ABIDEII-UCD | 32 | 43569 | 43565 | 14.58 (12, 17.83) | Siemens TriTim | 3T |
| ABIDEII-UCLA | 32 | 43480 | 43596 | 9.577 (7.751, 15.03) | Siemens TriTim | 3T |
| ABIDEII-USM | 33 | 43511 | 43537 | 19.72 (9.12, 38.86) | Siemens TriTim | 3T |
| Barcelona | 76 | 39/4 | 32/1 | 12.33 (7.17, 17.08) | Siemens Trio | 3T |
| BRC | 52 | 19/0 | 33/0 | 15 (10, 18) | GE Signa HDx | 3T |
| CMU | 27 | 43535 | 43534 | 27 (19, 40) | Siemens Magnetom | 3T |
| Dresden | 45 | 43542 | 43575 | 31.2 (21.1, 56.8) | Siemens Trio | 3T |
| FAIR | 85 | 36/7 | 27/15 | 11.56 (7.24, 15.94) | Siemens Magnetom | 3T |
| FRANKFURT | 27 | 43506 | 43509 | 18 (18, 18) | Siemens Sonata | 1.5T |
| FSM | 80 | 20/20 | 20/20 | 4.08 (1.83, 6) | GE Signa | 1.5T |
| MRC | 148 | 74/0 | 74/0 | 25 (18, 45) | GE Signa HDx | 3T |
| MYAD | 73 | 59/0 | 14/0 | 4.5 (1.5, 9) | Siemens symphony | 1.5T |
| NIJMEGEN1_1 | 33 | 43538 | 43510 | 15.09 (12.28, 18.02) | Siemens Trio | 3T |
| NIJMEGEN1_2* | 9 | - | 43473 | 17.47 (13.45, 18.52) | Siemens Trio | 3T |
| NIJMEGEN2 | 72 | 29/19 | 43693 | 26 (18, 40) | Siemens Avanto | 1.5T |
| NIJMEGEN3 | 95 | 37/4 | 45/9 | 9.71 (6.07, 12.27) | Siemens Avanto | 1.5T |
| ParelladaHGGM | 66 | 33/2 | 43495 | 13 (7, 18) | Philips Intera | 1.5T |
| PITT_1 | 56 | 43535 | 34/8 | 15 (8, 36) | Siemens Allegra | 3T |
| PITT_2 | 90 | 39/6 | 39/6 | 15 (8, 36) | Siemens Allegra | 3T |
| SAOPAULO | 35 | 15/0 | 20/0 | 11 (6, 19) | Philps | 3T |
| TCD_1 | 50 | 28/0 | 22/0 | 14.85 (10, 21.8) | Philips Achieva | 3T |
| TCD_2 | 39 | 17/0 | 22/0 | 15.9 (12, 25.9) | Philips Achieva | 3T |
| TORONTO_1 | 219 | 82/23 | 57/57 | 11.9 (3.3, 20.8) | Siemens Trio | 3T |
| TORONTO_2 | 203 | 107/43 | 29/24 | 11.05 (2.483, 21.67) | Siemens Trio | 3T |
| UMCU_1 | 88 | 41/7 | 37/3 | 12.06 (7.11, 24.72) | Philips | 1.5T |
| UMCU_2 | 9 | 6/0 | 43467 | 11.75 (9.96, 12.61) | Philips | 1.5T |
| **Total** | **3583** | **1778** | **1805** | | | |

* Excluded, as diagnosis would be completely confounded with random variable 'dataset' in the analysis model. For the calculation of the total scores (in **bold**) these datasets were also excluded.

**Table S2:** Association of rank-normalized ADOS severity scores with brain asymmetry

| | | $\beta$-value | | | Standard Error | | | $t$-value | | | $P$-value $^{\alpha}$ | | |
|---|---|---|---|---|---|---|---|---|---|---|---|---|---|
| AI region | N total | diag | age | sex | diag | age | sex | diag | age | sex | diag | age | sex |
| fusiform thickness | 1281 | 0.0013 | -0.0001 | 0.0028 | 0.0013 | 0.0001 | 0.0029 | 1.0471 | -0.9590 | 0.9919 | 0.2954 | 0.3379 | 0.3215 |
| inferiortemporal thickness | 1274 | 0.0005 | -0.0001 | 0.0009 | 0.0015 | 0.0001 | 0.0033 | 0.3475 | -0.3932 | 0.2647 | 0.7283 | 0.6942 | 0.7913 |
| isthmuscingulate thickness | 1281 | 0.0041 | 0.0005 | 0.0048 | 0.0016 | 0.0001 | 0.0043 | 2.5542 | 3.4803 | 1.1198 | **0.0108** | **0.0005** | 0.2631 |
| medialorbitofrontal thickness | 1296 | 0.0013 | 0.0003 | -0.0021 | 0.0018 | 0.0002 | 0.0041 | 0.7706 | 2.0702 | -0.5192 | 0.4412 | **0.0388** | 0.6037 |
| rostralanteriorcingulate thickness | 1285 | -0.0015 | -0.0001 | 0.0031 | 0.0026 | 0.0002 | 0.0057 | -0.5730 | -0.3898 | 0.5508 | 0.5668 | 0.6968 | 0.5820 |
| rostralmiddlefrontal thickness | 1269 | 0.0011 | -0.0003 | -0.0003 | 0.0012 | 0.0001 | 0.0026 | 0.8981 | -2.9926 | -0.1332 | 0.3694 | **0.0029** | 0.8940 |
| superiorfrontal thickness | 1274 | 0.0006 | -0.0000 | 0.0004 | 0.0009 | 0.0001 | 0.0020 | 0.6501 | -0.1801 | 0.2074 | 0.5158 | 0.8571 | 0.8358 |
| superiortemporal thickness | 1265 | -0.0002 | -0.0001 | 0.0007 | 0.0012 | 0.0001 | 0.0027 | -0.2116 | -0.6571 | 0.2434 | 0.8325 | 0.5113 | 0.8078 |
| total thickness | 1283 | -0.0005 | -0.0000 | -0.0008 | 0.0004 | 0.0000 | 0.0010 | -1.1316 | -0.7555 | -0.7904 | 0.2582 | 0.4502 | 0.4295 |
| medialorbitofrontalsurface area | 1280 | -0.0003 | 0.0003 | -0.0036 | 0.0028 | 0.0002 | 0.0063 | -0.1242 | 1.3555 | -0.5678 | 0.9012 | 0.1757 | 0.5704 |
| Putamen | 1242 | 0.0012 | 0.0003 | 0.0041 | 0.0014 | 0.0001 | 0.0032 | 0.8187 | 2.0149 | 1.2807 | 0.4132 | **0.0443** | 0.2007 |

$^{\alpha}$ Unadjusted, with in **bold** those values that are significant at the uncorrected level ($P < 0.05$).

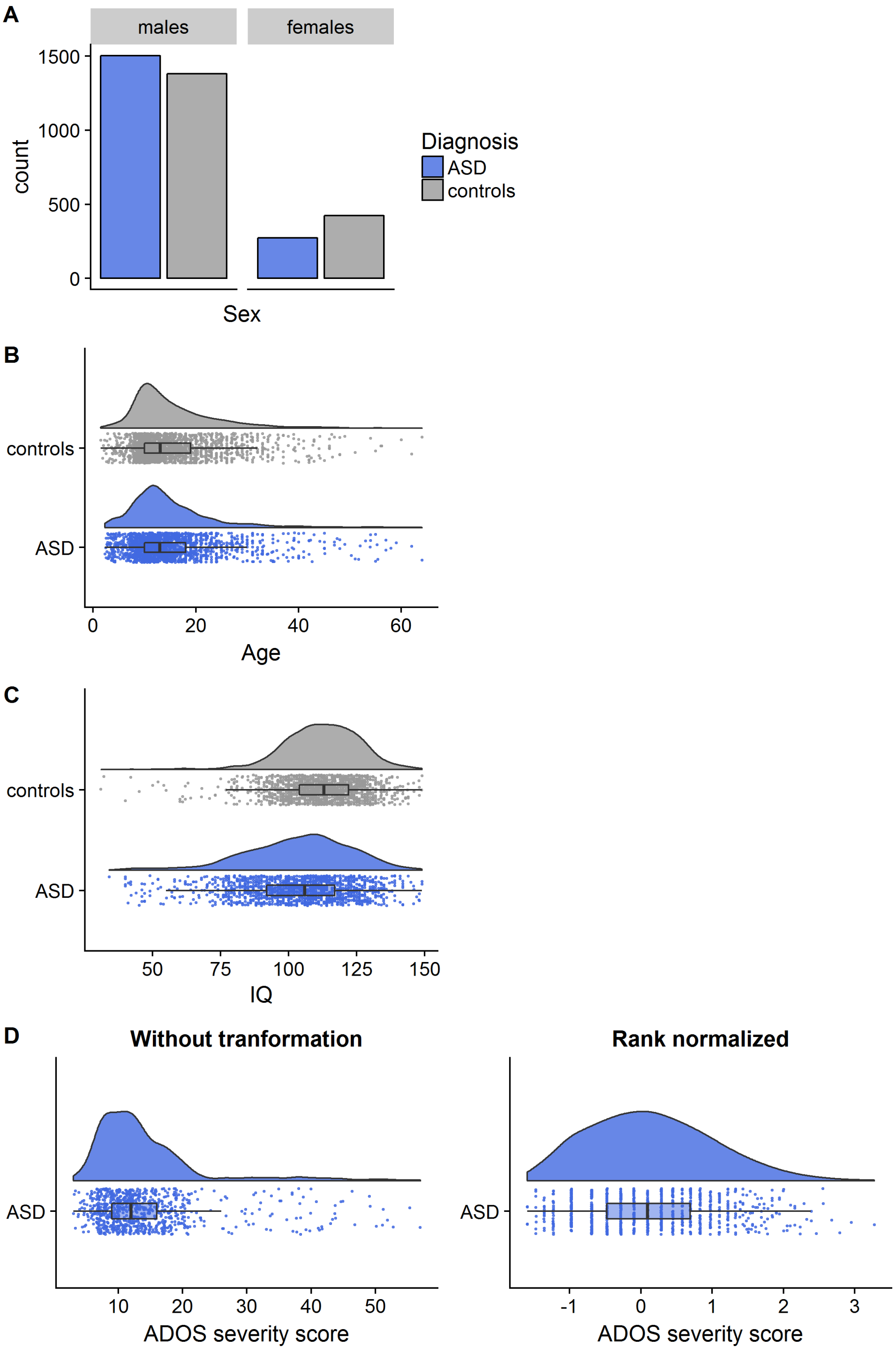

**Figure S1:** Distributions of (A) sex and (B) age in ASD and controls, and of (C) IQ and (D) ADOS severity score in ASD-only. The distributions of sex,age and IQ are comparable between the two diagnostic groups. ADOS severity score did not follow a normal distribution (left), so data were transformed using ranked normalization (right). *Blue* = ASD group; *grey* = control group.

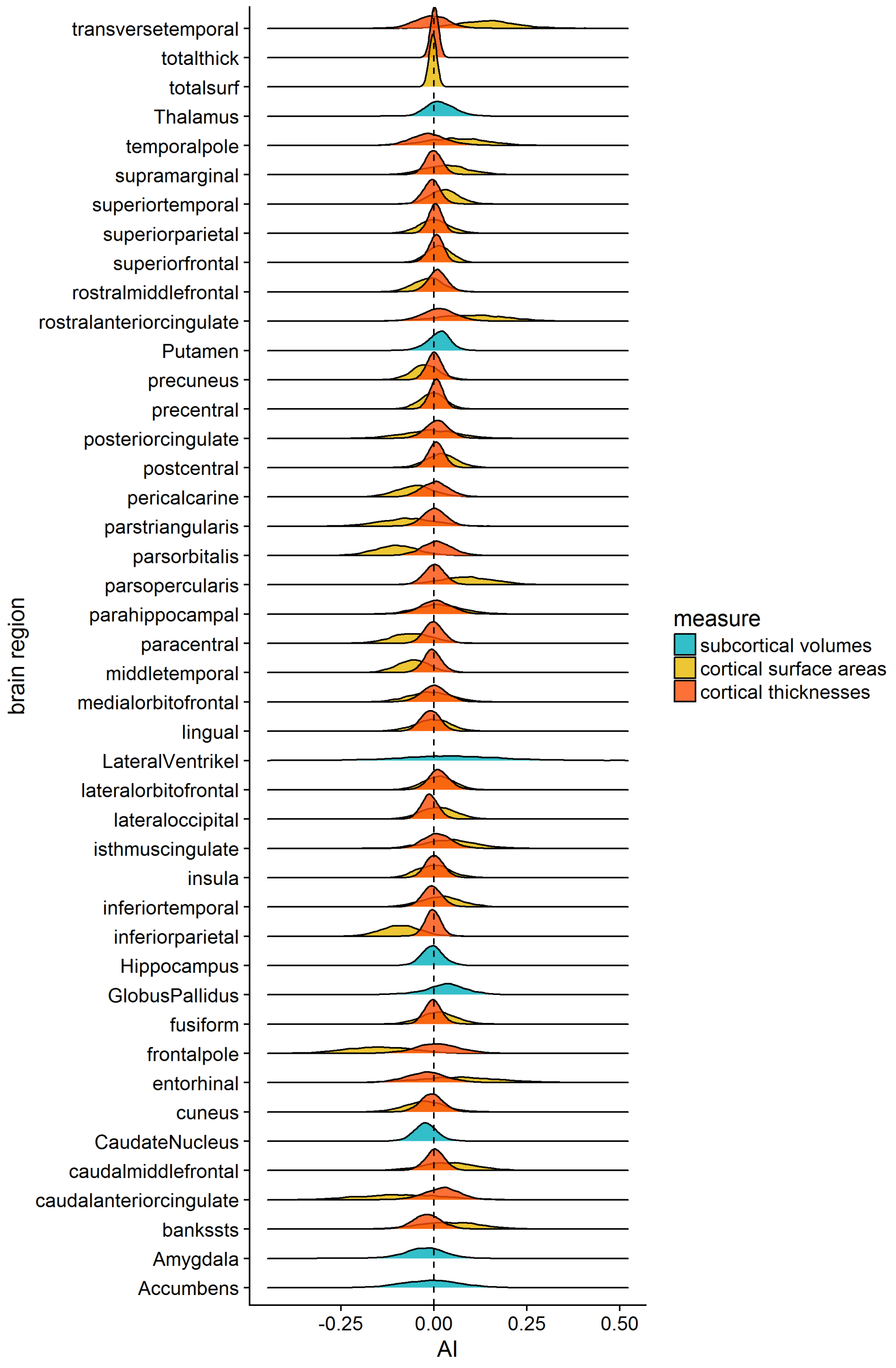

**Figure S2:** Joy plot of the distributions of AIs in the total sample. Shown for subcortical volumes (*cyan*), cortical surface areas (*orange*) and cortical thicknesses (*red*). Lateral ventricles were categorized among subcortical measures.

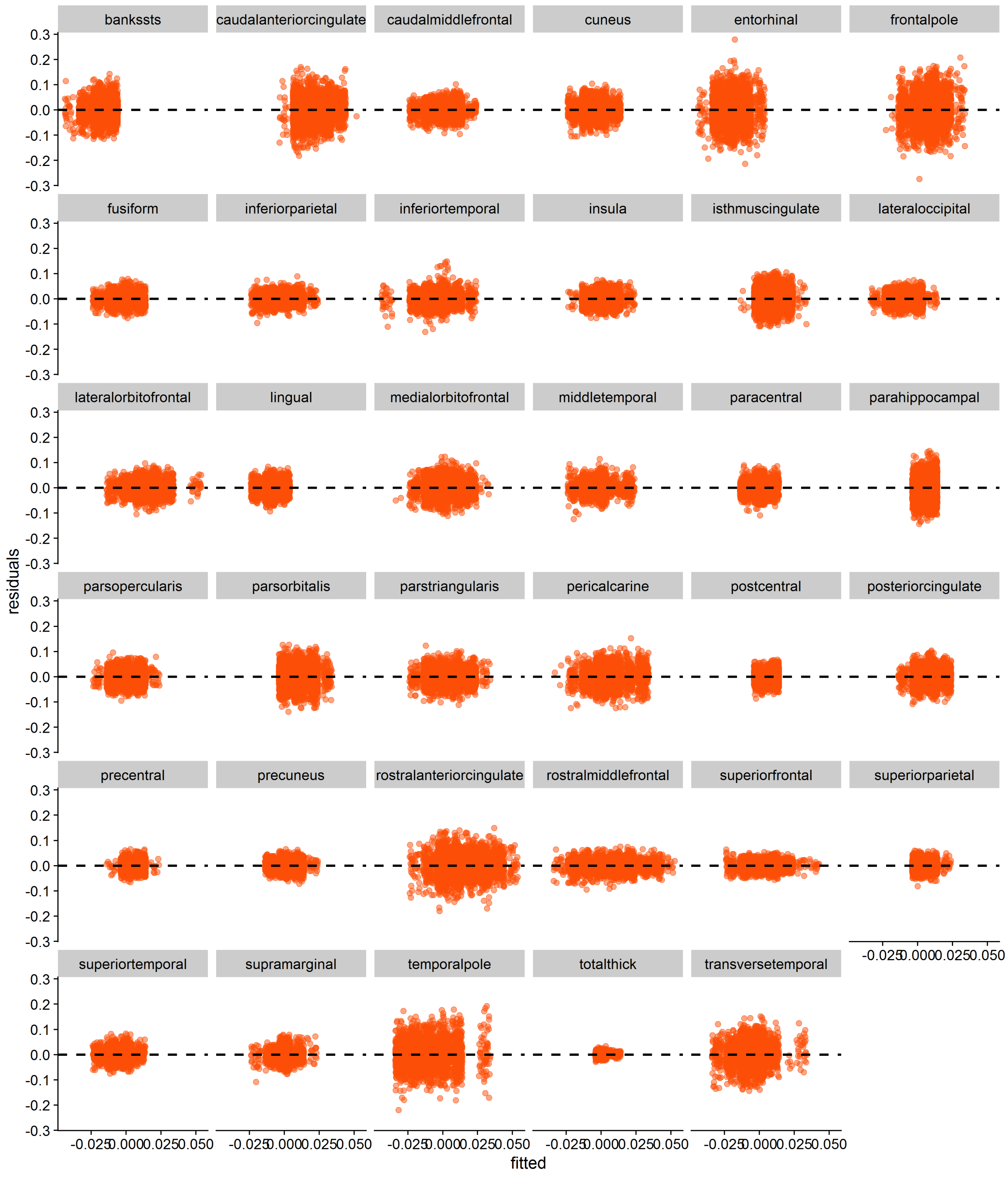

**Figure S3:** Residual plots of the linear mixed effects model analysis of cortical thickness AIs and the AI of the total average cortical thickness (totalthick).

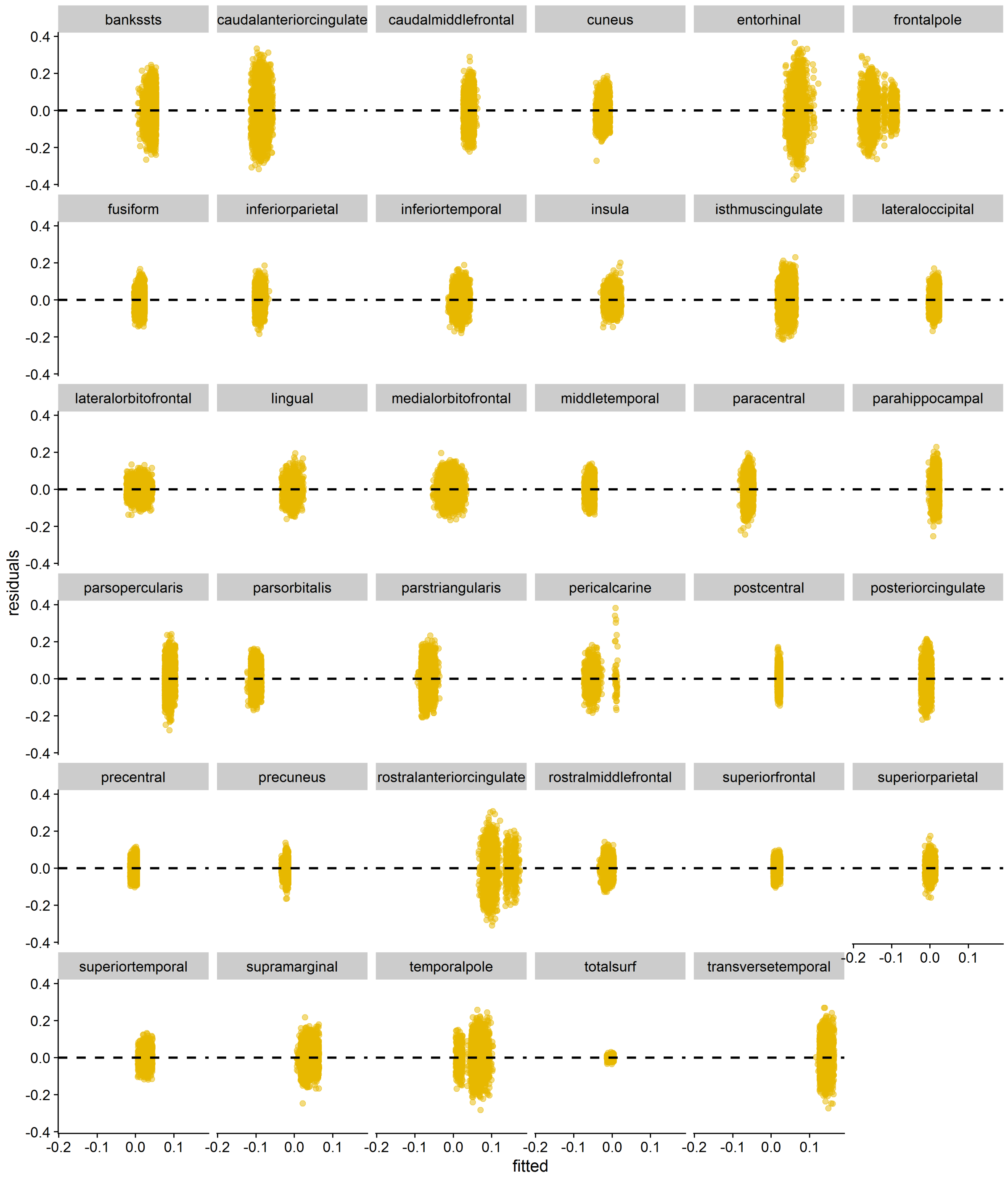

**Figure S4:** Residual plots of the linear mixed effects model analysis of cortical surface aera AIs and the AI of the total cortical surface area (totalsurf).

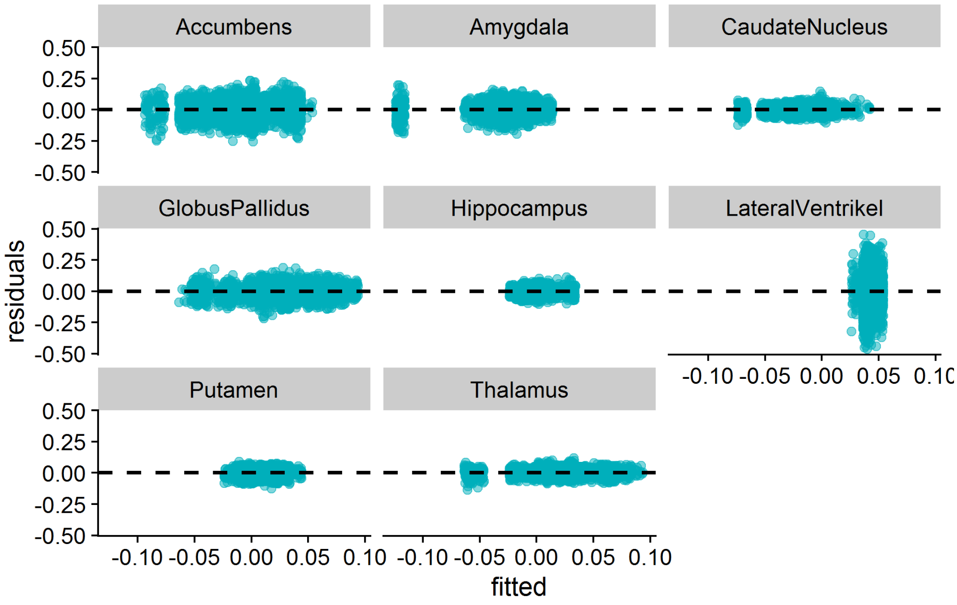

**Figure S5:** Residual plots from the linear mixed effects model analysis of subcortical volume AIs and the AI of the lateral ventricles.

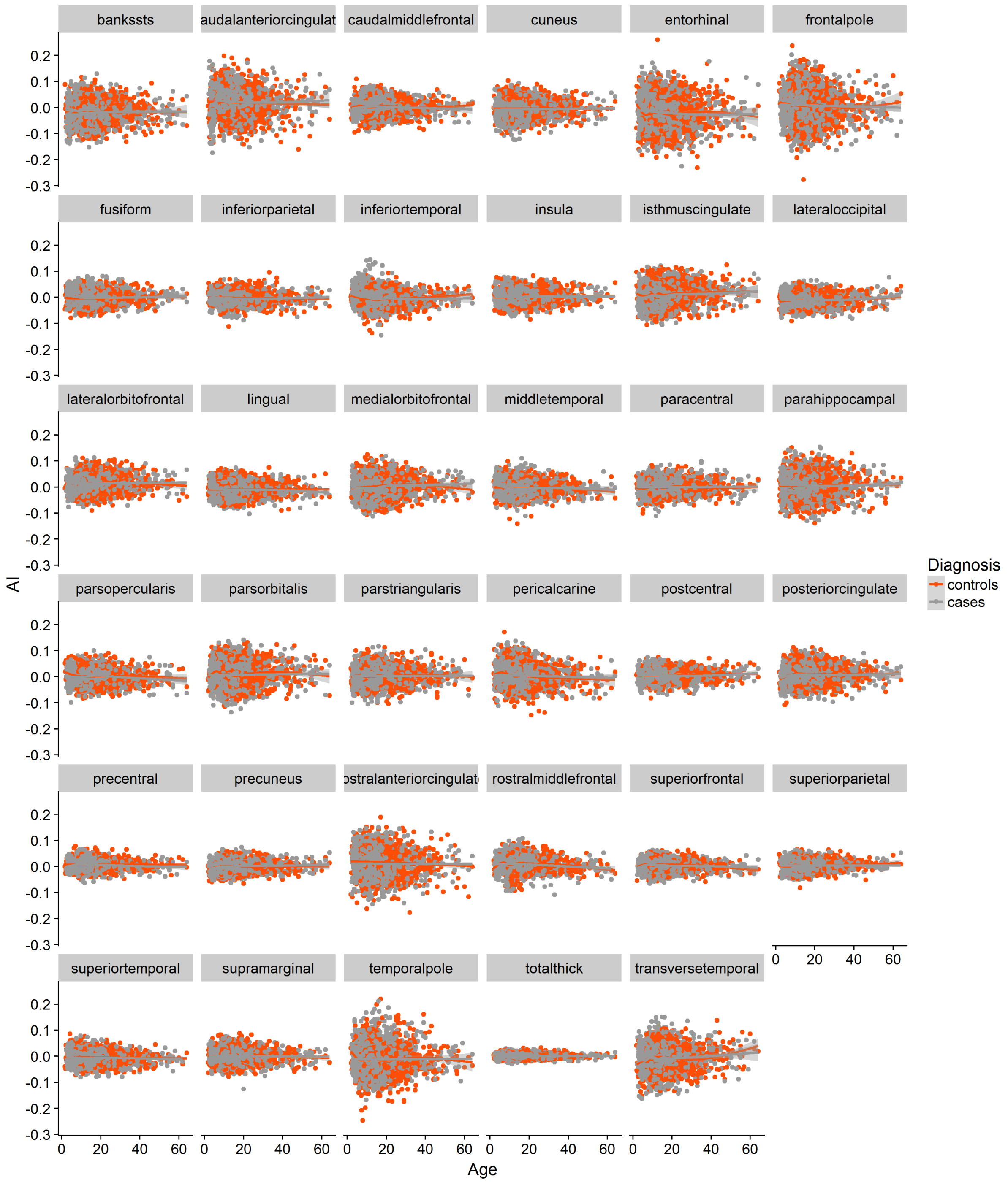

**Figure S6:** Scatter plots of the relationships between age and AIs of the total and regional cortical thicknesses.

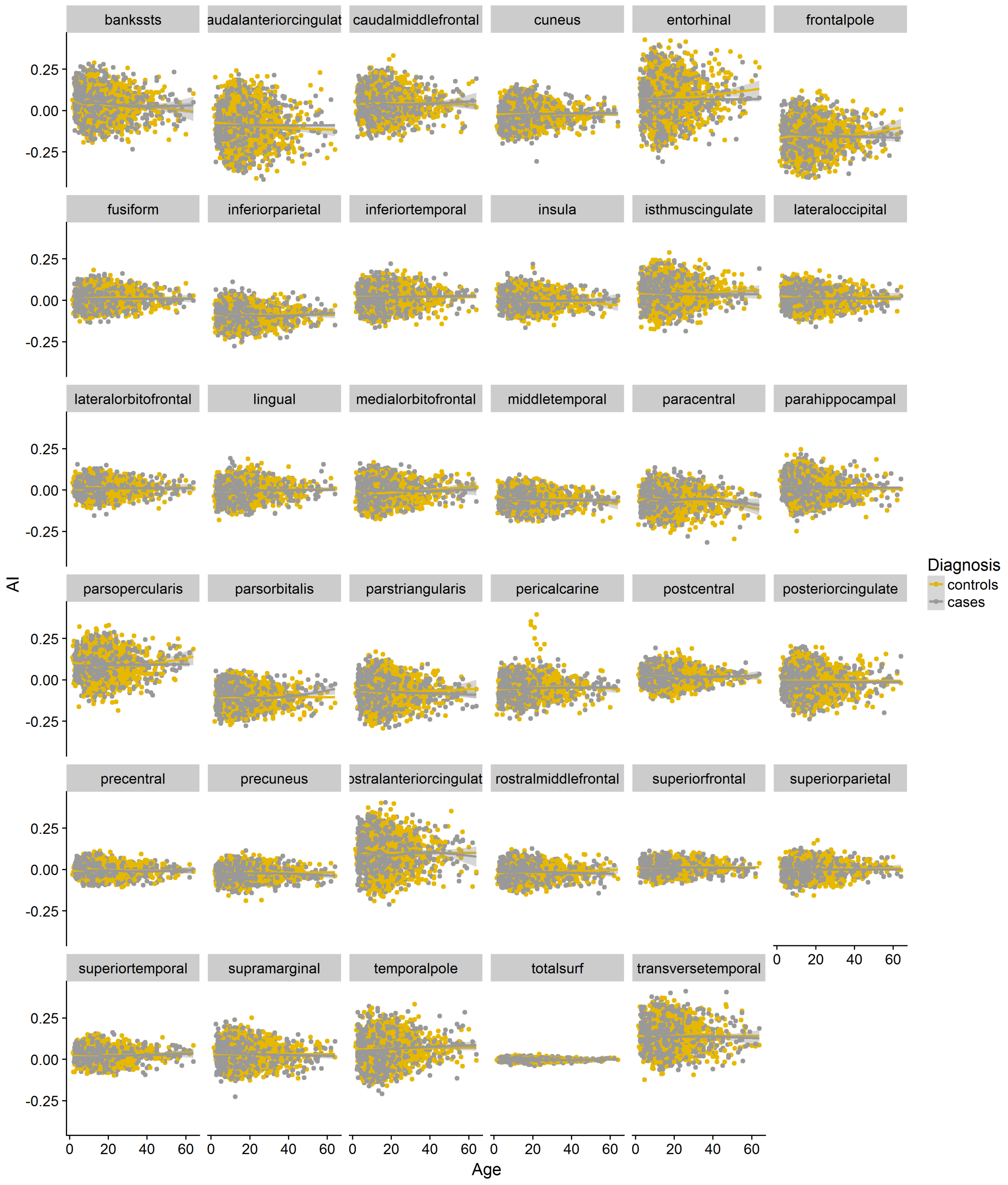

**Figure S7:** Scatter plots of the relationships between age and AIs of the total and regional cortical surface areas.

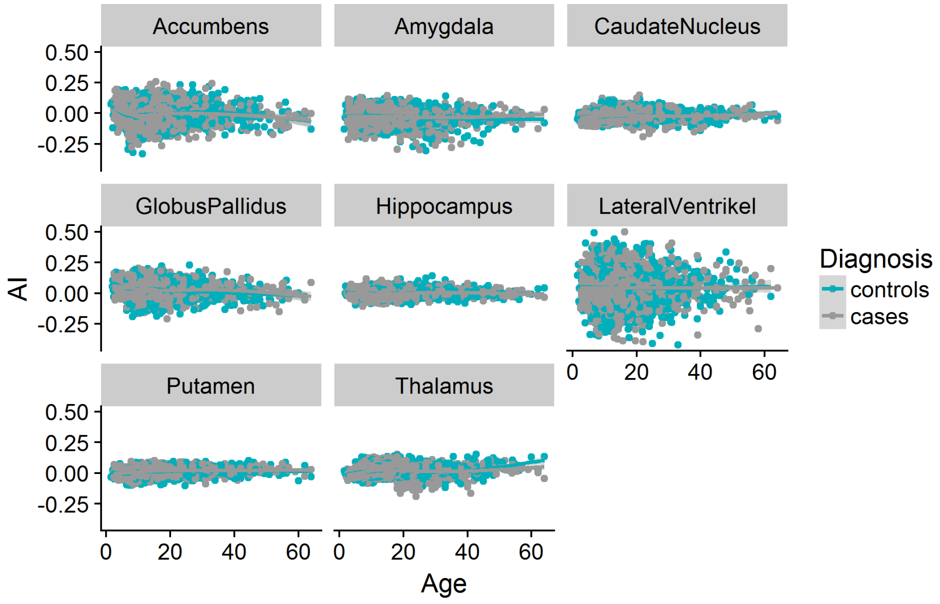

**Figure S8:** Scatter plots of the relationships between age and AIs of the subcortical volumes and lateral venctricles.

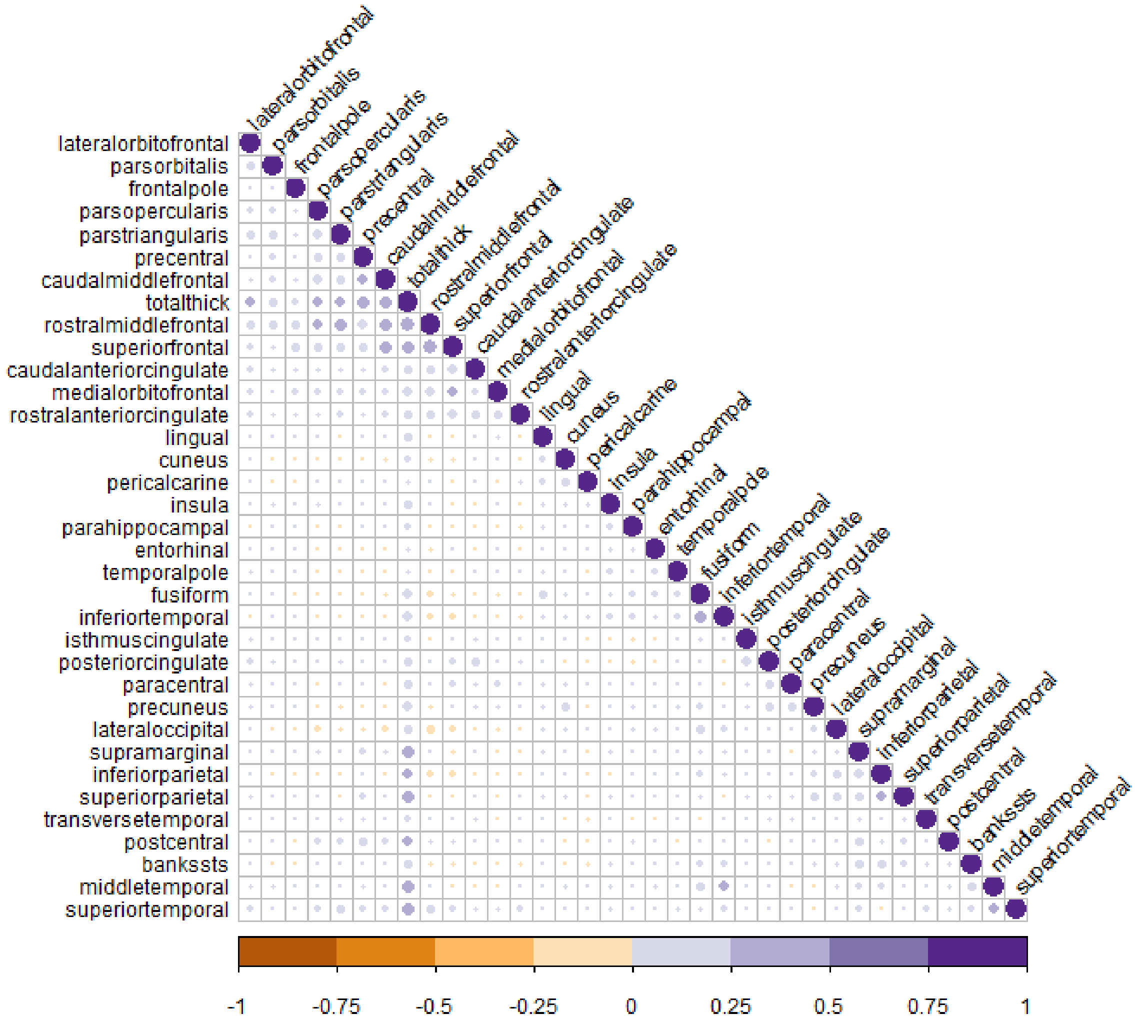

**Figure S9:** Correlations between AIs of cortical thickness. Correlations ranged from -0.15 to 0.47. Negative correlations are shown in orange, and positive correlations are shown in purple. Color intensities and circle sizes are proportional to the magnitudes of the correlation coefficients.

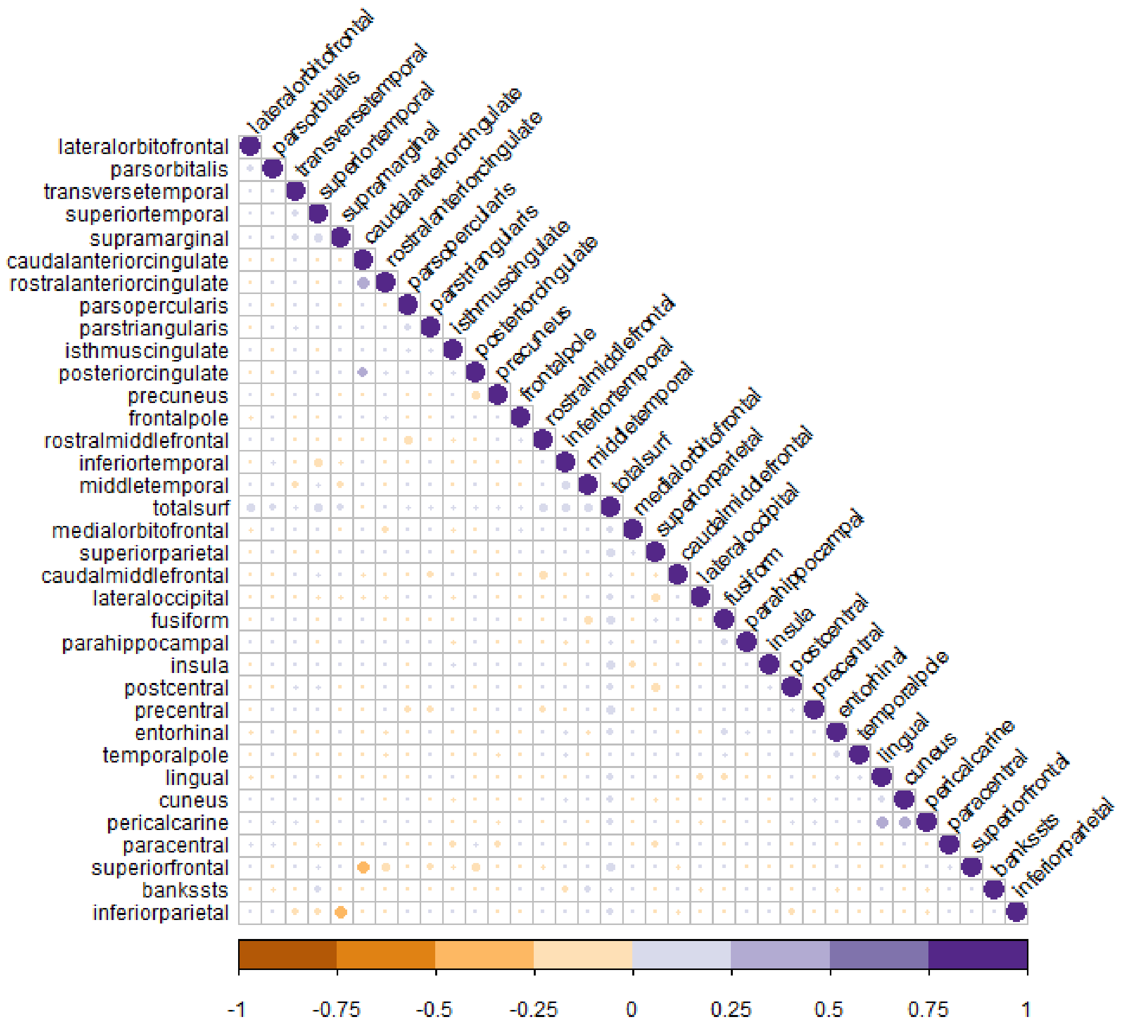

**Figure S10:** Correlations between AIs of cortical surface area. Correlations ranged from -0.37 to 0.36. Negative correlations are shown in orange, and positive correlations are shown in purple. Color intensities and circle sizes are proportional to the magnitudes of the correlation coefficients.

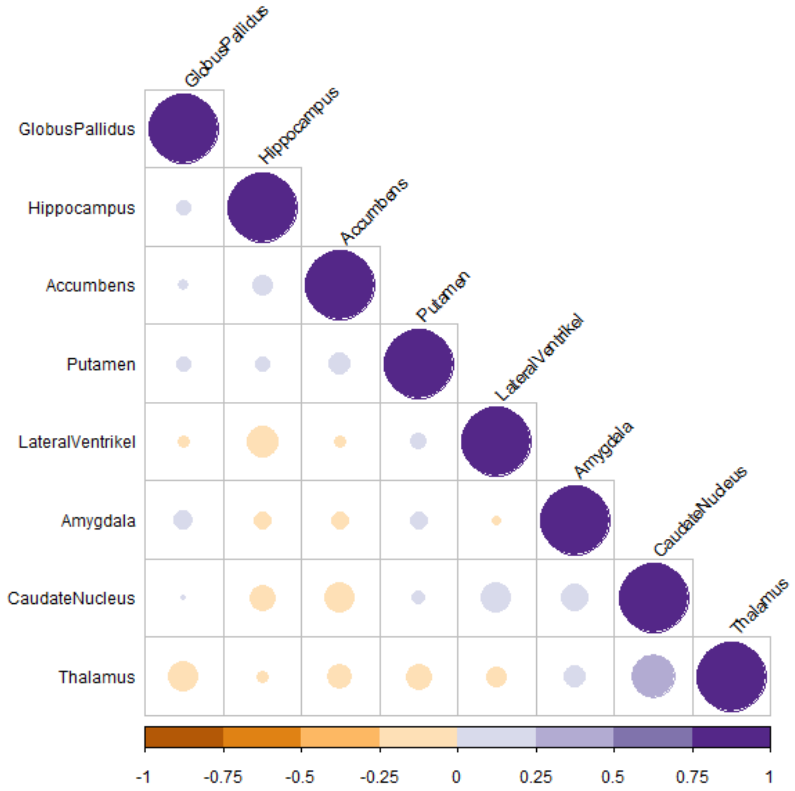

**Figure S11:** Correlations between AIs of subcortical volumes plus lateral ventricles. Correlations ranged from -0.37 to 0.47. Negative correlations are shown in orange, and positive correlations are shown in purple. Color intensities and circle sizes are proportional to the magnitudes of the correlation coefficients

**Table S3:** Linear mixed model results for cortical thickness AIs.

| | | $\beta$-value | | | Standard Error | | | $t$-value | | | $P$-value $^{\alpha}$ | | | |
|---|---|---|---|---|---|---|---|---|---|---|---|---|---|---|
| region | N cases/controls | diag | age | sex | diag | age | sex | diag | age | sex | diag | age | sex | Cohen's $d$ (95% CI) |
| bankssts | 1575/1648 | -0.0003 | -0.0000 | 0.0009 | 0.0014 | 0.0001 | 0.0018 | -0.1852 | -0.0625 | 0.4865 | 0.8531 | 0.9502 | 0.6267 | -0.007 (-0.08,0.06) |
| caudalanteriorcingulate | 1588/1654 | 0.0012 | 0.0000 | 0.0028 | 0.0017 | 0.0001 | 0.0022 | 0.7059 | 0.1288 | 1.2740 | 0.4803 | 0.8975 | 0.2028 | 0.025 (-0.04,0.09) |
| caudalmiddlefrontal | 1605/1654 | -0.0013 | -0.0000 | -0.0003 | 0.0009 | 0.0001 | 0.0012 | -1.4960 | -0.1545 | -0.2408 | 0.1348 | 0.8772 | 0.8097 | -0.053 (-0.12,0.02) |
| cuneus | 1602/1634 | -0.0021 | 0.0001 | -0.0010 | 0.0011 | 0.0001 | 0.0014 | -1.9467 | 1.3481 | -0.7058 | 0.0517 | 0.1777 | 0.4804 | -0.069 (-0.14,0) |
| entorhinal | 1593/1635 | 0.0025 | -0.0003 | -0.0028 | 0.0020 | 0.0001 | 0.0026 | 1.2768 | -2.0009 | -1.1040 | 0.2018 | **0.0455** | 0.2697 | 0.045 (-0.02,0.11) |
| frontalpole | 1617/1666 | -0.0006 | -0.0002 | -0.0028 | 0.0021 | 0.0002 | 0.0027 | -0.2682 | -1.6073 | -1.0274 | 0.7886 | 0.1081 | 0.3043 | -0.009 (-0.08,0.06) |
| fusiform | 1605/1637 | 0.0029 | -0.0000 | 0.0014 | 0.0008 | 0.0001 | 0.0010 | 3.7864 | -0.1845 | 1.3727 | **0.0002** | 0.8536 | 0.1700 | 0.134 (0.07,0.2) |
| inferiorparietal | 1589/1659 | -0.0016 | -0.0000 | -0.0002 | 0.0007 | 0.0001 | 0.0009 | -2.3278 | -0.3570 | -0.2571 | **0.0200** | 0.7211 | 0.7971 | -0.082 (-0.15,-0.01) |
| inferiortemporal | 1589/1646 | 0.0029 | -0.0001 | 0.0005 | 0.0010 | 0.0001 | 0.0013 | 3.0643 | -0.9891 | 0.3868 | **0.0022** | 0.3227 | 0.6989 | 0.109 (0.04,0.18) |
| insula | 1597/1654 | 0.0011 | 0.0001 | 0.0017 | 0.0008 | 0.0001 | 0.0011 | 1.3533 | 1.1434 | 1.5480 | 0.1760 | 0.2530 | 0.1217 | 0.048 (-0.02,0.12) |
| isthmuscingulate | 1594/1659 | -0.0046 | 0.0002 | 0.0007 | 0.0013 | 0.0001 | 0.0017 | -3.5204 | 2.0623 | 0.3892 | **0.0004** | **0.0393** | 0.6972 | -0.125 (-0.19,-0.06) |
| lateraloccipital | 1592/1659 | 0.0010 | 0.0001 | 0.0015 | 0.0008 | 0.0001 | 0.0010 | 1.3249 | 1.3464 | 1.4726 | 0.1853 | 0.1783 | 0.1409 | 0.047 (-0.02,0.12) |
| lateralorbitofrontal | 1609/1651 | 0.0001 | -0.0000 | -0.0011 | 0.0010 | 0.0001 | 0.0013 | 0.1192 | -0.1275 | -0.8438 | 0.9051 | 0.8986 | 0.3988 | 0.004 (-0.06,0.07) |
| lingual | 1614/1656 | -0.0005 | -0.0001 | -0.0032 | 0.0009 | 0.0001 | 0.0012 | -0.5740 | -0.8145 | -2.7515 | 0.5660 | 0.4154 | **0.0060** | -0.02 (-0.09,0.05) |
| medialorbitofrontal | 1626/1653 | -0.0044 | 0.0004 | -0.0007 | 0.0012 | 0.0001 | 0.0015 | -3.7654 | 3.9339 | -0.4827 | **0.0002** | **0.0001** | 0.6293 | -0.133 (-0.2,-0.06) |
| middletemporal | 1598/1624 | -0.0005 | 0.0000 | 0.0010 | 0.0009 | 0.0001 | 0.0011 | -0.5623 | 0.1292 | 0.8877 | 0.5740 | 0.8972 | 0.3747 | -0.02 (-0.09,0.05) |
| paracentral | 1604/1646 | 0.0009 | -0.0000 | -0.0007 | 0.0009 | 0.0001 | 0.0012 | 1.0059 | -0.0375 | -0.6095 | 0.3145 | 0.9701 | 0.5422 | 0.036 (-0.03,0.1) |
| parahippocampal | 1626/1677 | 0.0007 | 0.0001 | -0.0030 | 0.0015 | 0.0001 | 0.0019 | 0.5087 | 0.6148 | -1.5988 | 0.6110 | 0.5387 | 0.1100 | 0.018 (-0.05,0.09) |
| parsopercularis | 1553/1638 | -0.0014 | -0.0001 | 0.0017 | 0.0010 | 0.0001 | 0.0012 | -1.5004 | -1.0518 | 1.3666 | 0.1336 | 0.2930 | 0.1719 | -0.054 (-0.12,0.02) |
| parsorbitalis | 1596/1657 | -0.0026 | 0.0000 | 0.0003 | 0.0014 | 0.0001 | 0.0019 | -1.8359 | 0.2051 | 0.1484 | 0.0665 | 0.8375 | 0.8820 | -0.065 (-0.13,0) |
| parstriangularis | 1616/1661 | -0.0004 | -0.0001 | 0.0015 | 0.0011 | 0.0001 | 0.0014 | -0.3503 | -0.8275 | 1.0653 | 0.7262 | 0.4080 | 0.2868 | -0.012 (-0.08,0.06) |
| pericalcarine | 1566/1678 | -0.0003 | -0.0002 | -0.0000 | 0.0013 | 0.0001 | 0.0017 | -0.2143 | -2.0200 | -0.0184 | 0.8303 | **0.0435** | 0.9853 | -0.008 (-0.08,0.06) |
| postcentral | 1589/1628 | -0.0005 | 0.0001 | -0.0005 | 0.0007 | 0.0000 | 0.0010 | -0.7134 | 1.0843 | -0.4888 | 0.4757 | 0.2783 | 0.6251 | -0.025 (-0.09,0.04) |
| posteriorcingulate | 1607/1668 | -0.0016 | 0.0000 | -0.0001 | 0.0011 | 0.0001 | 0.0014 | -1.4507 | 0.1975 | -0.0705 | 0.1470 | 0.8435 | 0.9438 | -0.051 (-0.12,0.02) |
| precentral | 1548/1650 | -0.0010 | -0.0001 | -0.0006 | 0.0006 | 0.0000 | 0.0008 | -1.6628 | -2.2311 | -0.6827 | 0.0965 | **0.0257** | 0.4948 | -0.059 (-0.13,0.01) |
| precuneus | 1608/1671 | -0.0002 | 0.0000 | -0.0024 | 0.0007 | 0.0001 | 0.0009 | -0.3438 | 0.3077 | -2.7130 | 0.7310 | 0.7583 | **0.0067** | -0.012 (-0.08,0.06) |
| rostralanteriorcingulate | 1597/1655 | -0.0061 | -0.0001 | 0.0019 | 0.0016 | 0.0001 | 0.0021 | -3.9099 | -0.6723 | 0.9287 | **0.0001** | 0.5014 | 0.3531 | -0.138 (-0.21,-0.07) |
| rostralmiddlefrontal | 1597/1665 | -0.0030 | -0.0001 | 0.0010 | 0.0008 | 0.0001 | 0.0010 | -3.7997 | -1.9066 | 0.9986 | **0.0001** | 0.0567 | 0.3181 | -0.134 (-0.2,-0.07) |
| superiorfrontal | 1604/1650 | -0.0027 | 0.0000 | -0.0006 | 0.0006 | 0.0001 | 0.0008 | -4.5933 | 0.3614 | -0.7949 | **0.0000** | 0.7178 | 0.4267 | -0.162 (-0.23,-0.09) |
| superiorparietal | 1612/1670 | 0.0001 | 0.0000 | 0.0002 | 0.0006 | 0.0001 | 0.0008 | 0.1369 | 0.5740 | 0.2600 | 0.8911 | 0.5660 | 0.7949 | 0.005 (-0.06,0.07) |
| superiortemporal | 1573/1665 | -0.0020 | 0.0000 | 0.0017 | 0.0008 | 0.0001 | 0.0010 | -2.5121 | 0.4962 | 1.6251 | **0.0121** | 0.6198 | 0.1042 | -0.089 (-0.16,-0.02) |
| supramarginal | 1595/1669 | 0.0004 | -0.0001 | 0.0022 | 0.0008 | 0.0001 | 0.0010 | 0.5185 | -1.3799 | 2.1247 | 0.6042 | 0.1677 | 0.0337 | 0.018 (-0.05,0.09) |
| temporalpole | 1597/1630 | 0.0018 | 0.0000 | -0.0005 | 0.0018 | 0.0001 | 0.0024 | 1.0052 | 0.2665 | -0.2067 | 0.3149 | 0.7899 | 0.8362 | 0.036 (-0.03,0.1) |
| transversetemporal | 1615/1679 | 0.0006 | 0.0003 | 0.0021 | 0.0016 | 0.0001 | 0.0020 | 0.3670 | 2.6037 | 1.0045 | 0.7137 | **0.0093** | 0.3152 | 0.013 (-0.06,0.08) |
| totalthick | 1604/1669 | -0.0007 | -0.0000 | 0.0003 | 0.0003 | 0.0000 | 0.0003 | -2.5711 | -0.5710 | 0.7356 | **0.0102** | 0.5681 | 0.4620 | -0.091 (-0.16,-0.02) |

$^{\alpha}$ Unadjusted, with in**bold** those values that are significant at the uncorrected level ($P < 0.05$).

**Table S4:** Linear mixed model results for cortical surface area AIs.

| region | N cases/controls | $\beta$-value | | | Standard Error | | | $t$-value | | | $P$-value $^{\alpha}$ | | | Cohen's $d$ (95% CI) |
|---|---|---|---|---|---|---|---|---|---|---|---|---|---|---|
| | | diag | age | sex | diag | age | sex | diag | age | sex | diag | age | sex | |
| bankssts | 1587/1655 | 0.0036 | -0.0007 | -0.0009 | 0.0028 | 0.0002 | 0.0036 | 1.2625 | -4.2771 | -0.2524 | 0.2069 | **0.0000** | 0.8008 | 0.045 (-0.02,0.11) |
| caudalanteriorcingulate | 1606/1655 | -0.0026 | -0.0003 | 0.0083 | 0.0039 | 0.0003 | 0.0050 | -0.6704 | -1.2802 | 1.6472 | 0.5027 | 0.2006 | 0.0996 | -0.024 (-0.09,0.04) |
| caudalmiddlefrontal | 1594/1675 | -0.0046 | -0.0000 | 0.0062 | 0.0025 | 0.0002 | 0.0033 | -1.8302 | -0.0027 | 1.8990 | 0.0673 | 0.9978 | 0.0577 | -0.065 (-0.13,0) |
| cuneus | 1564/1633 | -0.0030 | 0.0000 | -0.0015 | 0.0019 | 0.0001 | 0.0025 | -1.5503 | 0.0684 | -0.5776 | 0.1212 | 0.9455 | 0.5636 | -0.055 (-0.12,0.01) |
| entorhinal | 1557/1617 | -0.0045 | 0.0006 | 0.0012 | 0.0039 | 0.0003 | 0.0051 | -1.1457 | 2.1699 | 0.2416 | 0.2520 | **0.0301** | 0.8091 | -0.041 (-0.11,0.03) |
| frontalpole | 1607/1663 | 0.0026 | 0.0002 | 0.0022 | 0.0030 | 0.0002 | 0.0039 | 0.8684 | 0.8238 | 0.5656 | 0.3852 | 0.4101 | 0.5717 | 0.031 (-0.04,0.1) |
| fusiform | 1596/1661 | -0.0038 | 0.0001 | 0.0019 | 0.0016 | 0.0001 | 0.0021 | -2.3200 | 0.9365 | 0.9113 | **0.0204** | 0.3491 | 0.3622 | -0.082 (-0.15,-0.01) |
| inferiorparietal | 1612/1668 | -0.0007 | -0.0001 | 0.0023 | 0.0018 | 0.0001 | 0.0023 | -0.3993 | -1.0181 | 0.9768 | 0.6897 | 0.3087 | 0.3287 | -0.014 (-0.08,0.05) |
| inferiortemporal | 1587/1655 | 0.0002 | 0.0001 | -0.0005 | 0.0019 | 0.0001 | 0.0025 | 0.0950 | 0.8456 | -0.2094 | 0.9243 | 0.3979 | 0.8342 | 0.003 (-0.07,0.07) |
| insula | 1599/1635 | 0.0021 | -0.0000 | -0.0022 | 0.0016 | 0.0001 | 0.0020 | 1.3251 | -0.3232 | -1.1054 | 0.1852 | 0.7466 | 0.2691 | 0.047 (-0.02,0.12) |
| isthmuscingulate | 1582/1663 | 0.0012 | 0.0000 | -0.0005 | 0.0024 | 0.0002 | 0.0031 | 0.4997 | 0.0669 | -0.1719 | 0.6173 | 0.9467 | 0.8635 | 0.018 (-0.05,0.09) |
| lateraloccipital | 1598/1656 | -0.0005 | -0.0001 | 0.0014 | 0.0017 | 0.0001 | 0.0021 | -0.3200 | -0.8750 | 0.6304 | 0.7490 | 0.3817 | 0.5285 | -0.011 (-0.08,0.06) |
| lateralorbitofrontal | 1599/1656 | -0.0038 | 0.0001 | -0.0000 | 0.0014 | 0.0001 | 0.0018 | -2.7132 | 0.7323 | -0.0182 | **0.0067** | 0.4641 | 0.9855 | -0.096 (-0.16,-0.03) |
| lingual | 1570/1615 | 0.0020 | 0.0001 | 0.0013 | 0.0017 | 0.0001 | 0.0022 | 1.1445 | 1.1455 | 0.5683 | 0.2525 | 0.2521 | 0.5699 | 0.041 (-0.03,0.11) |
| medialorbitofrontal | 1608/1663 | 0.0079 | 0.0002 | -0.0028 | 0.0019 | 0.0002 | 0.0025 | 4.1539 | 1.0264 | -1.1226 | **0.0000** | 0.3048 | 0.2617 | 0.147 (0.08,0.22) |
| middletemporal | 1600/1671 | 0.0009 | -0.0001 | 0.0008 | 0.0015 | 0.0001 | 0.0020 | 0.5850 | -1.0012 | 0.3989 | 0.5586 | 0.3168 | 0.6900 | 0.021 (-0.05,0.09) |
| paracentral | 1575/1635 | 0.0015 | -0.0002 | -0.0012 | 0.0020 | 0.0001 | 0.0026 | 0.7479 | -1.8108 | -0.4547 | 0.4546 | 0.0703 | 0.6494 | 0.027 (-0.04,0.1) |
| parahippocampal | 1580/1603 | 0.0000 | -0.0002 | 0.0004 | 0.0022 | 0.0001 | 0.0028 | 0.0205 | -1.4337 | 0.1577 | 0.9836 | 0.1517 | 0.8747 | 0.001 (-0.07,0.07) |
| parsopercularis | 1558/1648 | -0.0003 | 0.0002 | -0.0003 | 0.0026 | 0.0002 | 0.0034 | -0.1089 | 1.1364 | -0.0980 | 0.9133 | 0.2559 | 0.9219 | -0.004 (-0.07,0.07) |
| parsorbitalis | 1595/1645 | 0.0045 | 0.0001 | 0.0016 | 0.0020 | 0.0001 | 0.0025 | 2.2896 | 0.5529 | 0.6486 | **0.0221** | 0.5804 | 0.5166 | 0.081 (0.01,0.15) |
| parstriangularis | 1612/1669 | 0.0006 | -0.0002 | 0.0032 | 0.0026 | 0.0002 | 0.0033 | 0.2440 | -1.0387 | 0.9771 | 0.8072 | 0.2990 | 0.3286 | 0.009 (-0.06,0.08) |
| pericalcarine | 1574/1630 | 0.0008 | -0.0000 | -0.0011 | 0.0019 | 0.0002 | 0.0025 | 0.4335 | -0.2025 | -0.4238 | 0.6647 | 0.8395 | 0.6717 | 0.015 (-0.05,0.08) |
| postcentral | 1586/1629 | -0.0000 | 0.0000 | 0.0005 | 0.0015 | 0.0001 | 0.0019 | -0.0173 | 0.1423 | 0.2565 | 0.9862 | 0.8869 | 0.7976 | -0.001 (-0.07,0.07) |
| posteriorcingulate | 1603/1666 | -0.0051 | -0.0002 | 0.0031 | 0.0025 | 0.0002 | 0.0032 | -2.0516 | -1.0265 | 0.9683 | **0.0403** | 0.3047 | 0.3330 | -0.072 (-0.14,0) |
| precentral | 1579/1637 | -0.0030 | -0.0002 | -0.0009 | 0.0012 | 0.0001 | 0.0015 | -2.4560 | -2.1227 | -0.5822 | **0.0141** | **0.0339** | 0.5605 | -0.087 (-0.16,-0.02) |
| precuneus | 1582/1634 | -0.0013 | 0.0001 | 0.0002 | 0.0013 | 0.0001 | 0.0017 | -0.9729 | 1.0797 | 0.1241 | 0.3307 | 0.2803 | 0.9013 | -0.035 (-0.1,0.03) |
| rostralanteriorcingulate | 1612/1685 | -0.0033 | 0.0001 | 0.0043 | 0.0033 | 0.0002 | 0.0043 | -0.9990 | 0.2507 | 1.0184 | 0.3178 | 0.8021 | 0.3086 | -0.035 (-0.1,0.03) |
| rostralmiddlefrontal | 1610/1660 | 0.0017 | -0.0000 | -0.0003 | 0.0014 | 0.0001 | 0.0019 | 1.2128 | -0.0172 | -0.1415 | 0.2253 | 0.9863 | 0.8874 | 0.043 (-0.03,0.11) |
| superiorfrontal | 1600/1679 | 0.0013 | -0.0001 | -0.0008 | 0.0011 | 0.0001 | 0.0015 | 1.1434 | -0.8963 | -0.5521 | 0.2530 | 0.3701 | 0.5809 | 0.04 (-0.03,0.11) |
| superiorparietal | 1589/1655 | 0.0002 | 0.0000 | -0.0025 | 0.0015 | 0.0001 | 0.0019 | 0.1213 | 0.2385 | -1.3477 | 0.9035 | 0.8115 | 0.1778 | 0.004 (-0.06,0.07) |
| superiortemporal | 1592/1670 | 0.0005 | 0.0001 | 0.0001 | 0.0014 | 0.0001 | 0.0018 | 0.3974 | 0.6498 | 0.0823 | 0.6911 | 0.5158 | 0.9344 | 0.014 (-0.05,0.08) |
| supramarginal | 1594/1685 | -0.0000 | -0.0000 | -0.0058 | 0.0021 | 0.0002 | 0.0027 | -0.0042 | -0.2396 | -2.1091 | 0.9967 | 0.8106 | **0.0350** | 0 (-0.07,0.07) |
| temporalpole | 1585/1652 | -0.0008 | 0.0003 | 0.0026 | 0.0027 | 0.0002 | 0.0036 | -0.3062 | 1.2256 | 0.7265 | 0.7595 | 0.2204 | 0.4676 | -0.011 (-0.08,0.06) |
| transversetemporal | 1628/1672 | 0.0009 | -0.0003 | 0.0011 | 0.0027 | 0.0002 | 0.0034 | 0.3450 | -1.4585 | 0.3147 | 0.7301 | 0.1448 | 0.7530 | 0.012 (-0.06,0.08) |
| totalsurf | 1588/1649 | 0.0001 | -0.0000 | 0.0003 | 0.0002 | 0.0000 | 0.0003 | 0.5623 | -0.9166 | 0.7997 | 0.5740 | 0.3594 | 0.4239 | 0.02 (-0.05,0.09) |

$^{\alpha}$ Unadjusted, with in**bold** those values that are significant at the uncorrected level ($P < 0.05$).

**Table S5:** Linear mixed model results for subcortical volume AIs.

| | | $\beta$-value | | | Standard Error | | | *t*-value | | | *P*-value $^{\alpha}$ | | | |
|---|---|---|---|---|---|---|---|---|---|---|---|---|---|---|
| region | N cases/controls | diag | age | sex | diag | age | sex | diag | age | sex | diag | age | sex | Cohen's *d* (95% CI) |
| Accumbens | 1631/1666 | -0.0034 | -0.0003 | 0.0018 | 0.0026 | 0.0002 | 0.0034 | -1.3108 | -1.2784 | 0.5120 | 0.1900 | 0.2012 | 0.6087 | -0.046 (-0.11,0.02) |
| Amygdala | 1588/1660 | -0.0017 | 0.0000 | 0.0043 | 0.0019 | 0.0002 | 0.0024 | -0.9161 | 0.2262 | 1.7631 | 0.3597 | 0.8210 | 0.0780 | -0.032 (-0.1,0.04) |
| CaudateNucleus | 1580/1655 | 0.0008 | 0.0001 | 0.0034 | 0.0010 | 0.0001 | 0.0013 | 0.7811 | 0.9118 | 2.6328 | 0.4348 | 0.3619 | **0.0085** | 0.028 (-0.04,0.1) |
| GlobusPallidus | 1570/1636 | 0.0027 | -0.0006 | 0.0008 | 0.0019 | 0.0002 | 0.0024 | 1.4650 | -3.4441 | 0.3093 | 0.1430 | **0.0006** | 0.7571 | 0.052 (-0.02,0.12) |
| Hippocampus | 1555/1614 | -0.0014 | -0.0001 | -0.0010 | 0.0010 | 0.0001 | 0.0013 | -1.3904 | -1.1497 | -0.7301 | 0.1645 | 0.2503 | 0.4654 | -0.05 (-0.12,0.02) |
| LateralVentrikel | 1518/1577 | 0.0001 | -0.0002 | 0.0077 | 0.0049 | 0.0003 | 0.0061 | 0.0194 | -0.8609 | 1.2600 | 0.9845 | 0.3893 | 0.2078 | 0.001 (-0.07,0.07) |
| Putamen | 1559/1675 | 0.0035 | 0.0003 | 0.0014 | 0.0010 | 0.0001 | 0.0013 | 3.5347 | 3.2742 | 1.0633 | **0.0004** | **0.0011** | 0.2877 | 0.126 (0.06,0.19) |
| Thalamus | 1599/1647 | 0.0012 | 0.0005 | 0.0025 | 0.0010 | 0.0001 | 0.0014 | 1.1525 | 4.9237 | 1.8477 | 0.2492 | **0.0000** | 0.0647 | 0.041 (-0.03,0.11) |

$^{\alpha}$ Unadjusted, with in**bold** those values that are significant at the uncorrected level ($P < 0.05$).

**Table S6:** Linear mixed model results after no outliers were removed, after a non-linear effect of age was added to the model, and after the two youngest datasets were removed.

| | no outliers | | non-linear age | | no youngest datasets | |
|---|---|---|---|---|---|---|
| region | $P^{\alpha}$ | $d^{\beta}$ | $P^{\alpha}$ | $d^{\beta}$ | $P^{\alpha}$ | $d^{\beta}$ |
| fusiform thickness | **0.0014** | 0.1095 | **0.0002** | 0.1336 | **0.0005** | 0.1267 |
| inferiortemporal thickness | **0.0030** | 0.1016 | **0.0026** | 0.1069 | **0.0052** | 0.1016 |
| isthmuscingulate thickness | *0.0098* | -0.0885 | **0.0004** | -0.1246 | **0.0010** | -0.1189 |
| medialorbitofrontal thickness | **0.0005** | -0.1188 | **0.0002** | -0.1326 | **0.0001** | -0.1421 |
| rostralanteriorcingulate thickness | **0.0008** | -0.1155 | **0.0001** | -0.1382 | **0.0000** | -0.1526 |
| rostralmiddlefrontal thickness | **0.0033** | -0.1006 | **0.0002** | -0.1337 | **0.0000** | -0.1544 |
| superiorfrontal thickness | **0.0001** | -0.1342 | **0.0000** | -0.1619 | **0.0000** | -0.1706 |
| superiortemporal thickness | *0.0490* | -0.0676 | **0.0122** | -0.0890 | *0.0149* | -0.0885 |
| total thickness | *0.0504* | -0.0670 | **0.0103** | -0.0906 | **0.0074** | -0.0968 |
| medialorbitofrontalsurface area | *0.0015* | 0.1085 | **0.0000** | 0.1459 | **0.0000** | 0.1486 |
| Putamen | **0.0007** | 0.1162 | **0.0004** | 0.1270 | **0.0003** | 0.1311 |

$^{\alpha}$ Unadjusted *P*-value, related to the effect of diagnosis. Values in **bold** are significant at FDR ¡ 0.05, and those in *italic* are significant at the uncorrected level ($P < 0.05$). $^{\beta}$ Cohen's *d* effect size, related to the effect of diagnoiss.

**Table S7:** Directions of asymmetry changes in cases versus controls.

| | mean ± SD | | Cohen's *d* (95% CI) | | Direction of asymmetry | |
|---|---|---|---|---|---|---|
| Region | AI controls | AI cases | Left | Right | controls | cases |
| fusiform thickness | -0.004 ± 0.02 | -0.001 ± 0.02 | -0.201 (-0.27,-0.13) | -0.285 (-0.35,-0.22) | rightward | decreased |
| inferiortemporal thickness | -0.006 ± 0.03 | -0.003 ± 0.03 | -0.143 (-0.21,-0.07) | -0.218 (-0.29,-0.15) | rightward | decreased |
| isthmuscingulate thickness | 0.012 ± 0.04 | 0.006 ± 0.04 | -0.027 (-0.1,0.04) | 0.091 (0.02,0.16) | leftward | decreased |
| medialorbitofrontal thickness | 0.003 ± 0.03 | -0.001 ± 0.03 | 0.014 (-0.05,0.08) | 0.121 (0.05,0.19) | leftward | reversed |
| rostralanteriorcingulate thickness | 0.016 ± 0.04 | 0.011 ± 0.05 | -0.128 (-0.2,-0.06) | 0.017 (-0.05,0.09) | leftward | decreased |
| rostralmiddlefrontal thickness | 0.01 ± 0.03 | 0.008 ± 0.03 | -0.018 (-0.09,0.05) | 0.061 (-0.01,0.13) | leftward | decreased |
| superiorfrontal thickness | 0.006 ± 0.02 | 0.005 ± 0.02 | -0.069 (-0.14,0) | 0.018 (-0.05,0.09) | leftward | decreased |
| superiortemporal thickness | -0.005 ± 0.02 | -0.007 ± 0.02 | -0.13 (-0.2,-0.06) | -0.065 (-0.13,0) | rightward | increased |
| total thickness | 0.002 ± 0.01 | 0.001 ± 0.01 | -0.148 (-0.22,-0.08) | -0.111 (-0.18,-0.04) | leftward | decreased |
| medialorbitofrontalsurface area | -0.014 ± 0.06 | -0.005 ± 0.06 | 0.094 (0.03,0.16) | -0.019 (-0.09,0.05) | rightward | decreased |
| Putamen | 0.012 ± 0.03 | 0.013 ± 0.03 | -0.022 (-0.09,0.05) | -0.084 (-0.15,-0.01) | leftward | increased |

The mean and standard deviation of the AI in controls is indicated for each AI which showed a significant effect of diagnosis in the primary analysis. Also the Cohen's d effect sizes for left and right hemispheric measures are indicated, as well as the direction of asymmetry in controls (derived from the average AI) and its change in cases. Positive AI values indicate leftward asymmetry, negative AI values indicate rightward asymmetry.

**Table S8:** Age:diagnosis and sex:diagnosis interaction effects.

| | Age | | | Sex | | |
|---|---|---|---|---|---|---|
| region | age:diag | children | adults | sex:diag | males | females |
| fusiform thickness | 0.2786 | | | 0.5021 | | |
| inferiortemporal thickness | 0.4839 | | | 0.5866 | | |
| isthmuscingulate thickness | 0.8322 | | | 0.0659 | | |
| medialorbitofrontal thickness | 0.3997 | | | 0.3847 | | |
| rostralanteriorcingulate thickness | 0.6900 | | | **0.0011** | $<$ **0.001** | 0.224 |
| rostralmiddlefrontal thickness | 0.8211 | | | 0.9539 | | |
| superiorfrontal thickness | 0.1478 | | | 0.0732 | | |
| superiortemporal thickness | 0.1327 | | | 0.1680 | | |
| total thickness | 0.5103 | | | 0.1406 | | |
| medialorbitofrontalsurface area | 0.9504 | | | 0.7384 | | |
| putamen | 0.1822 | | | 0.6397 | | |

*P*-values for the interaction effects on AIs are indicated, for all AIs which showed significant effects of diagnosis in the primary analysis. Furthermore, *P*-values for the effects of diagnosis on AIs are indicated when tested separately in subsets: children, adults, males and females.

**Table S9:** Association of IQ with brain asymmetry within cases.

| | | $\beta$-value | | | Standard Error | | | $t$-value | | | $P$-value $^{\alpha}$ | | |
|---|---|---|---|---|---|---|---|---|---|---|---|---|---|
| AI region | N total | IQ | age | sex | IQ | age | sex | IQ | age | sex | IQ | age | sex |
| fusiform thickness | 1281 | -0.0000 | -0.0000 | 0.0009 | 0.0000 | 0.0001 | 0.0017 | -1.2842 | -0.0917 | 0.4884 | 0.1993 | 0.9270 | 0.6254 |
| inferiortemporal thickness | 1274 | -0.0001 | -0.0001 | -0.0015 | 0.0000 | 0.0001 | 0.0022 | -1.3452 | -0.6759 | -0.6772 | 0.1788 | 0.4992 | 0.4984 |
| isthmuscingulate thickness | 1281 | -0.0001 | 0.0003 | 0.0052 | 0.0001 | 0.0001 | 0.0029 | -1.0921 | 2.2581 | 1.8223 | 0.2750 | **0.0241** | 0.0687 |
| medialorbitofrontal thickness | 1296 | -0.0000 | 0.0004 | -0.0003 | 0.0001 | 0.0001 | 0.0026 | -0.2733 | 2.9299 | -0.1287 | 0.7847 | **0.0035** | 0.8976 |
| rostralanteriorcingulate thickness | 1285 | 0.0002 | -0.0000 | 0.0083 | 0.0001 | 0.0002 | 0.0035 | 3.3327 | -0.2226 | 2.3540 | **0.0009** | 0.8239 | **0.0187** |
| rostralmiddlefrontal thickness | 1269 | -0.0000 | -0.0002 | 0.0004 | 0.0000 | 0.0001 | 0.0017 | -0.9161 | -2.4466 | 0.2362 | 0.3598 | **0.0146** | 0.8133 |
| superiorfrontal thickness | 1274 | -0.0000 | 0.0000 | 0.0005 | 0.0000 | 0.0001 | 0.0013 | -0.0556 | 0.4243 | 0.3621 | 0.9556 | 0.6714 | 0.7174 |
| superiortemporal thickness | 1265 | 0.0000 | -0.0000 | 0.0006 | 0.0000 | 0.0001 | 0.0018 | 0.5866 | -0.3131 | 0.3455 | 0.5576 | 0.7542 | 0.7298 |
| total thickness | 1283 | -0.0000 | -0.0000 | -0.0005 | 0.0000 | 0.0000 | 0.0006 | -0.4332 | -0.6154 | -0.8282 | 0.6650 | 0.5384 | 0.4077 |
| medialorbitofrontalsurface area | 1280 | -0.0001 | 0.0003 | -0.0009 | 0.0001 | 0.0002 | 0.0042 | -0.7544 | 1.4024 | -0.2264 | 0.4508 | 0.1610 | 0.8210 |
| Putamen | 1242 | -0.0000 | 0.0002 | 0.0036 | 0.0000 | 0.0001 | 0.0022 | -0.4594 | 1.3817 | 1.6616 | 0.6460 | 0.1673 | 0.0969 |

$^{\alpha}$ Unadjusted, with in**bold** those values that are significant at the uncorrected level ($P < 0.05$).
